# Dual symbiosis in the deep-sea hydrothermal vent snail *Gigantopelta aegis* revealed by its hologenome

**DOI:** 10.1101/2020.09.23.308304

**Authors:** Yi Lan, Jin Sun, Chong Chen, Yanan Sun, Yadong Zhou, Yi Yang, Weipeng Zhang, Runsheng Li, Kun Zhou, Wai Chuen Wong, Yick Hang Kwan, Aifang Cheng, Salim Bougouffa, Cindy Lee Van Dover, Jian-Wen Qiu, Pei-Yuan Qian

**Affiliations:** Department of Ocean Science, Division of Life Science and Hong Kong Branch of the Southern Marine Science and Engineering Guangdong Laboratory (Guangzhou), The Hong Kong University of Science and Technology, Hong Kong, China; X-STAR, Japan Agency for Marine-Earth Science and Technology (JAMSTEC), 2-15 Natsushima-cho, Yokosuka, Kanagawa Prefecture 237-0061, Japan; Key Laboratory of Marine Ecosystem Dynamics, Second Institute of Oceanography, Ministry of Natural Resources, Hangzhou, China; College of Marine Life Science, Ocean University of China, 5 Yushan Road, Qingdao, 266003, China; Department of Biology, Hong Kong Baptist University, Kowloon Tong, Hong Kong; Computational Bioscience Research Centre, King Abdullah University of Science and Technology, Thuwal, 23955-6900, Saudi Arabia; King Abdullah University of Science and Technology (KAUST), Core Labs, Thuwal, 23955-6900, Saudi Arabia; Division of Marine Science and Conservation, Nicholas School of the Environment, Duke University, Beaufort, NC, United States

**Keywords:** cryptometamorphosis, methane-oxidising, sulphur-oxidising, Mollusca

## Abstract

Animals endemic to deep-sea hydrothermal vents often form obligatory relationships with bacterial symbionts, maintained by intricate host-symbiont interactions. Endosymbiosis with more than one symbiont is uncommon, and most genomic studies focusing on such ‘dual symbiosis’ systems have not investigated the host and the symbionts to a similar depth simultaneously. Here, we report a novel dual symbiosis among the peltospirid snail *Gigantopelta aegis* and its two Gammaproteobacteria endosymbionts – one being a sulphur oxidiser and the other a methane oxidiser. We assembled high-quality genomes for all three parties of this holobiont, with a chromosome-level assembly for the snail host (1.15 Gb, N50 = 82 Mb, 15 pseudo-chromosomes). In-depth analyses of these genomes reveal an intimate mutualistic relationship with complementarity in nutrition and metabolic codependency, resulting in a system highly versatile in transportation and utilisation of chemical energy. Moreover, *G. aegis* has an enhanced immune capability that likely facilitates the possession of more than one type of symbiont. Comparisons with *Chrysomallon squamiferum*, another chemosymbiotic snail in the same family but only with one sulphur-oxidising endosymbiont, show that the two snails’ sulphur-oxidising endosymbionts are phylogenetically distant, agreeing with previous results that the two snails have evolved endosymbiosis independently and convergently. Notably, the same capabilities of biosynthesis of specific nutrition lacking in the host genome are shared by the two sulphur-oxidising endosymbionts of the two snail genera, which may be a key criterion in the selection of symbionts by the hosts.

## Introduction

Animals live in harmony with microorganisms in and around them, forming ecological units referred to as holobionts. Some metazoans, for instance reef-forming corals, have established intricate obligatory symbiotic relationships with single-celled organisms that enable the holobiont to more effectively utilise the available energy source around them^1^. In deep-sea chemosynthetic environments such as hydrothermal vents, many endemic animals harbour endosymbiotic bacteria within their cells in order to directly access chemosynthesis utilising reducing substances such as hydrogen sulphide, hydrogen, thiosulfate, and methane^2^. Most of the species have evolved long-term symbiotic relationships with one dominant phylotype of microbial endosymbiont^3,4,5,6^. Several species of annelids and molluscs, however, are capable of hosting multiple co-occurring endosymbionts^7,8,9,10^. A number of studies have used genomic tools to explore the molecular mechanisms that enable and support symbioses^3,4,11^, but most studies focused more closely on either the symbiont or the host. Studying the symbiotic relationship by a total ‘hologenome’ approach combining high-quality genomes of both the host and the symbiont is key to understanding how the partners cooperate and collaborate to adapt and flourish in extreme environments.

Among deep-sea vent molluscs housing endosymbionts, the two peltospirid genera *Gigantopelta* and *Chrysomallon* are perhaps the most extreme ‘holobionts’. They host the symbionts inside a specialised organ within the gut (a modified oesophageal gland) rather than on the gill epithelium like all others^4,8,12,13,14,15^. This means that their endosymbionts are unable to exchange material directly with the vent fluid and are completely reliant on the host for supplies of reducing substances. The two genera are thought to have evolved this symbiosis independently through convergent evolution^15^. A crucial difference between the two is that while *Chrysomallon* relies on endosymbionts for nutrition throughout its post-settlement life^3,14^, *Gigantopelta* initially adopts grazing for nutrients and only later (at around 5-7 mm shell length) rapidly shifts to hosting endosymbionts^16^. This process, dubbed ‘cryptometamorphosis’, occurs with a dramatic reorganisation of its digestive system where the oesophageal gland is greatly enlarged while the other parts of the digestive system effectively stop growing^16^.

The fact that representatives of the two genera, *Gigantopelta aegis* and the Scaly-foot Snail *Chrysomallon squamiferum*, co-occur in great abundance in the Longqi vent field on the ultra-slow spreading Southwest Indian Ridge (SWIR) makes them ideal candidates to compare molecular adaptations in two similar-yet-different holobionts. From previous studies, we know that *C. squamiferum* hosts only one phylotype of sulphur-oxidising Gammaproteobacteria whose genome has been sequenced^3^. The whole genome of the host was also recently sequenced at chromosome-level^17^, but little has been explored in terms of its adaptations related to symbiosis. In the present study, we reveal that *Gigantopelta aegis* has a dual endosymbiosis, where the snail houses two Gammaproteobacteria endosymbionts, one sulphur-oxidising and one methane-oxidising, in its oesophageal gland. Previously, the 16S rRNA sequencing of the closely related *Gigantopelta chessoia* only revealed a single sulphur-oxidising Gammaproteobacteria symbiont^18^, although the transmission electron micrograph of the oesophageal gland hinted a second, rarer, methane-oxidising symbiont^15^. Here, we use high-quality genome assemblies of both the *G. aegis* snail host and its two physiologically distinct endosymbionts to elucidate interactions and co-operations among this ‘trinity’ of parties in the holobiont. Furthermore, we compare these results to the interactions between *C. squamiferum* and its single sulphur-oxidising endosymbiont in order to understand how may *Gigantopelta aegis* benefit from symbiosis from more than one symbiont.

## Results and Discussion

### A Holobiont with Three Parties and Hologenome Features

The two genera, *Gigantopelta aegis* (dark orange) and the Scaly-foot Snail *Chrysomallon squamiferum* (black), occur in great abundance side-by-side at Tiamat chimney in the Longqi vent field (**Fig. 1a**). We obtained high-quality genome assemblies of all three parties in the *G. aegis* holobiont using an adult snail 4 cm in shell length, for in-depth analyses of its symbiosis. The host genome had a size of 1.15 Gb, comprising 15 pseudo-chromosomes with an N50 of ∼82 Mb (93.8% genome completeness; **Fig. 1b, Table S1**). This is one of only a few chromosome-level genome assemblies currently available in Gastropoda or even Mollusca. The genome size of *G. aegis* is more than double of the *C. squamiferum* genome (∼444.4 Mb)^17^ mainly due to the larger contribution of repetitive regions (**Fig. S1, Fig. S2, Table S2**). The genomes of both *G. aegis* and *C. squamiferum* snails had 11 *hox* gene clusters in the same order (**Fig. S3**), 13 protein-coding genes of mito-genome in the same order (**Fig. S4**), and one-to-one chromosomal synteny (**Fig. 1c**). These indicate that much of the gene order and synteny are conserved within the family Peltospiridae, and such clear correspondences exemplify the high quality of the genome assembly for both genomes (**Fig. S5**). In addition, the genomic rearrangements that were marked by gene synteny were exclusively restricted to intra-chromosomal rather than inter-chromosomal (**Fig. 1c**). This result is also in line with our former comparative genomic analysis among four congener apple snail genomes, and the intra-chromosomal genomic rearrangements were more likely to be dominant than inter-chromosomal in shaping the genomic differences in Mollusca^19^. The genetic regulatory networks and related functional differences among them are underpinned by intra-chromosomal rearrangements, but how exactly warrants further studies. A total of 21,472 genes (**Table S3**) were predicted from the *G. aegis* genome, of which 1,783 genes were highly expressed in the oesophageal gland. No particular chromosomal distribution bias was observed among these genes (**Fig. 1b**).

**Figure 1.**
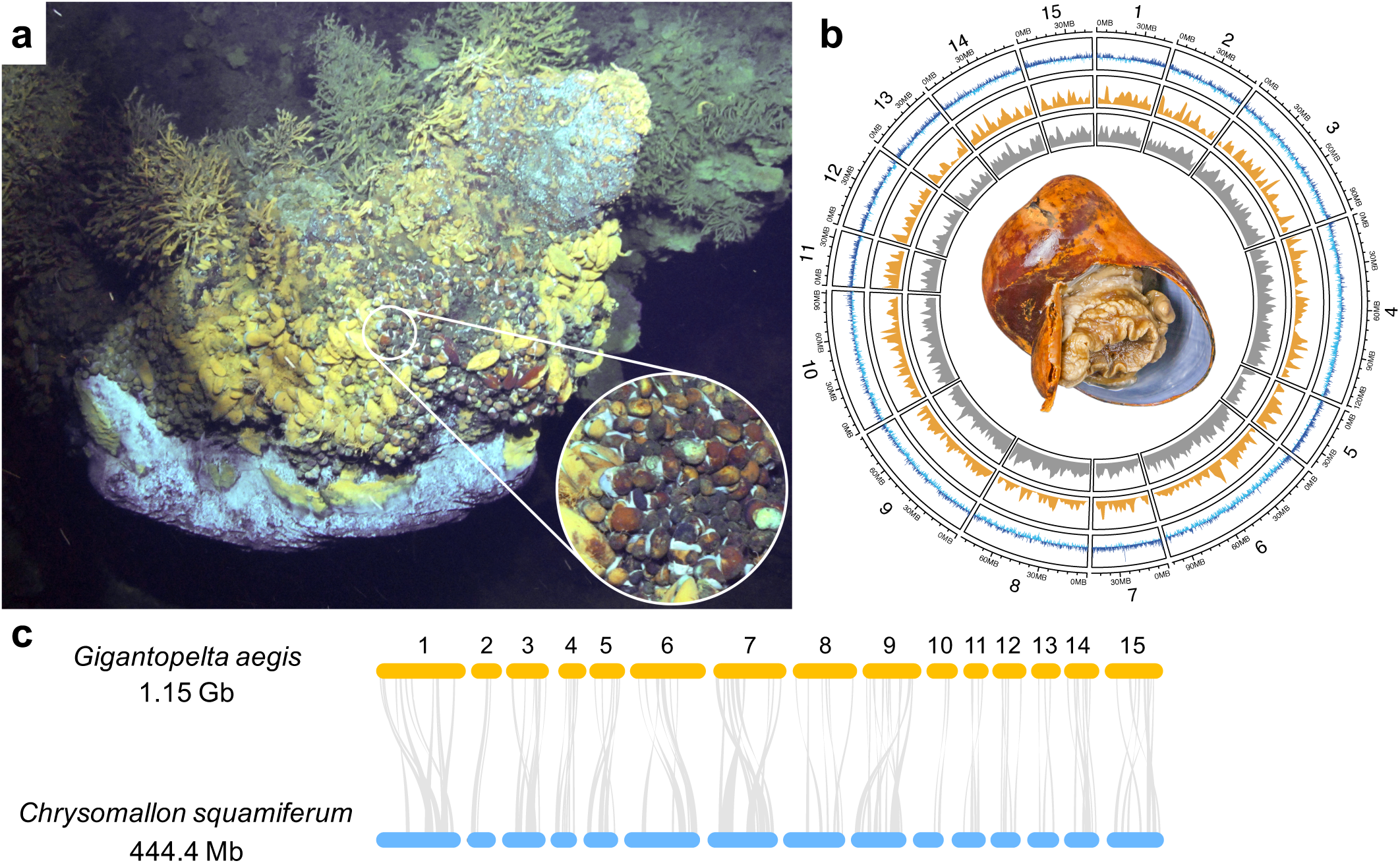
**a**. Two chemosymbiotic peltospirid snails, *Gigantopelta aegis* and the Scaly-foot Snail *Chrysomallon squamiferum*, occuring in great abundance side-by-side at Tiamat chimney in the Longqi vent field on the ultra-slow spreading Southwest Indian Ridge (*G. aegis* in dark orange; *C. squamiferum* in black). **b**. A circos plot shows key features of 15 pseudo-chromosomal linkage groups of the *Gigantopelta aegis* genome showing an adult snail at the centre, the inner circle (gray) indicating the gene density of each pseudo-chromosome, the second circle (orange) indicating the gene density of highly expressed genes in the oesophageal gland (*n* = 4), and the outer circle (blue) showing the GC content of the deviation from the average 37.22%. A sliding window of 100 kb in a step of 50kb was applied in the calculation of the GC content. **c**. Synteny blocks shared between *G. aegis* (orange) and *C. squamiferum* (blue). The pseudo-chromosome numbers of both *G. aegis* and *C. squamiferum* are 15. The synteny blocks are distributed in one-to-one correspondence among the pseudo-chromosomes without cross-chromosomal synteny.

Transmission electron micrographs of the oesophageal gland from *Gigantopelta aegis* confirmed the existence of intracellular endosymbionts densely packed inside bacteriocytes (**Fig. 2a, Fig. S6**). Two types of endosymbionts with distinct morphology features were found within a single bacteriocyte (**Fig. 2a**). Sequences of the 16S ribosomal RNA from oesophageal glands of three *G. aegis* individuals revealed that the two types of symbionts both belong to Gammaproteobacteria, the more abundant one being sulphur-oxidising bacteria (SOB hereafter) and the rarer one being methane-oxidising bacteria (MOB hereafter) in the family Methylococcaceae. Indeed, of the two symbionts identified from TEM, one exhibited intracellular stacked membranes (**Fig. 2a**) which is a characteristic of Type I methanotroph^12^, and the other lacked it. Fluorescence *in situ* hybridisation (FISH) carried out on transverse sections of the oesophageal gland from *G. aegis* yielded positive signals of both the SOB and the MOB (**Fig. 2b, Fig. S6**). These results confirm that *G. aegis* is a holobiont with three active parties, including the host and two physiologically distinct types of endosymbionts from Gammaproteobacteria. Genome binning results showed the SOB, the MOB, and the host could be separated with high fidelity by their GC content and sequencing coverage (**Fig. S7**). Both the TEM results and the sequencing coverage indicate that both endosymbionts are common in the oesophageal gland, with the SOB being approximate seven times more abundant than the MOB in this individual (**Table S4**). However, the relative abundance of the two symbionts possibly varies due to the fluctuant concentration of sulphur substances and methane, as was observed in *Bathymodiolus* mussels which have two symbionts^20,21^.

**Figure 2.**
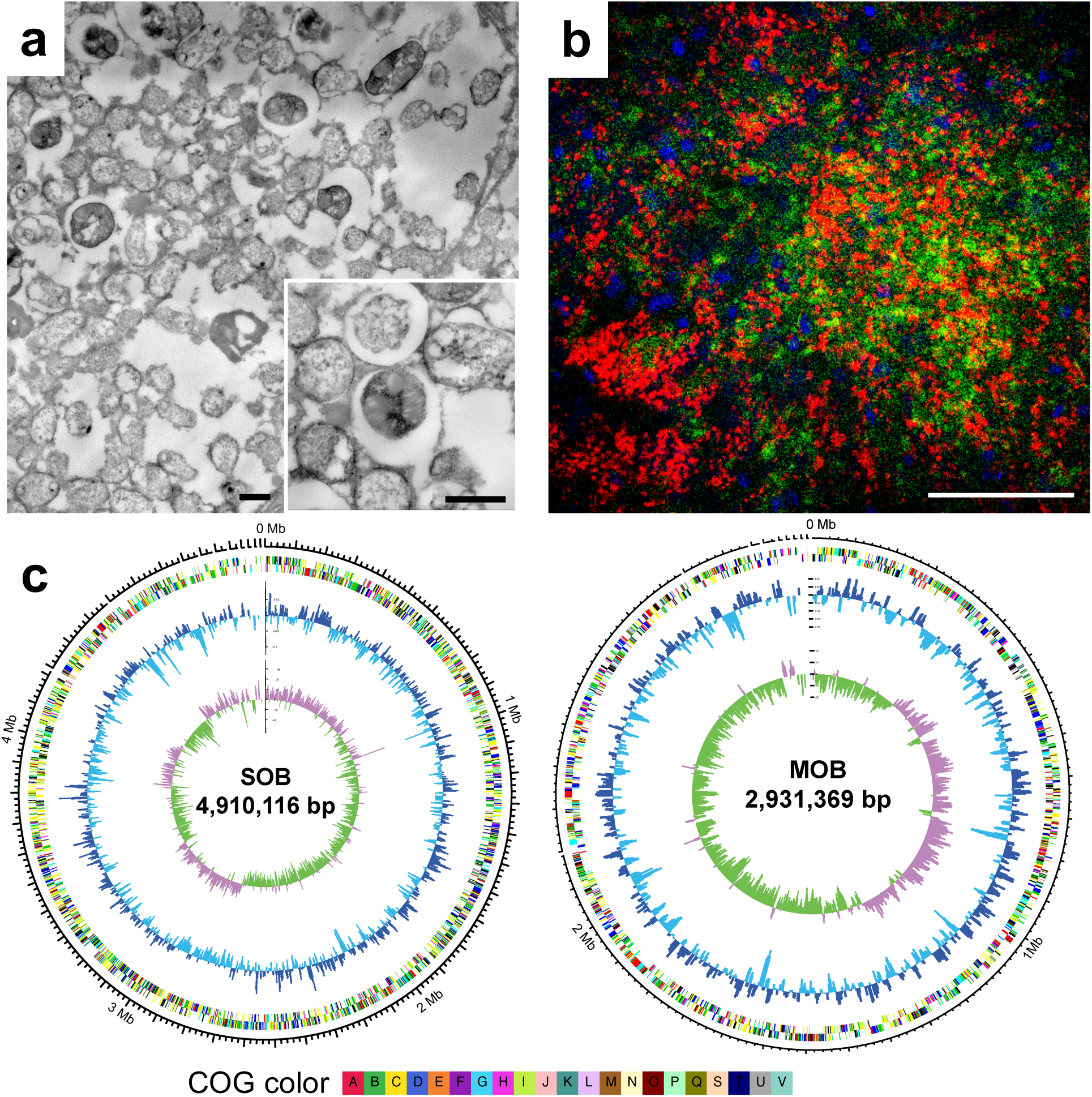
**a**. Transmission electron microscopy (TEM) images of intracellular endosymbionts showing two distinct morphological types, including one endosymbiont with intracellular stacked membranes and another without (scale bar: 1 µm). **b**. Fluorescence *in situ* hybridisation (FISH) image yielding signals of sulphur-oxidising endosymbionts (SOB; Cy3: green) and the methane-oxidising endosymbionts (MOB; Cy5: red) on transverse sections of the oesophageal gland from *Gigantopelta aegis* (scale bar: 50 µm). Host nuclear DNA with DAPI staining is blue. More detailed TEM and FISH images are shown in Figure S6. **c**. Key genome features of both the SOB and the MOB of *G. aegis* with the inner circle showing the GC skew, the second circle showing the GC content of the deviation from the average (SOB: 39.93%; MOB: 45.26%), and the outer circle showing the COG annotation categories with different colours. A sliding window of 5 kb in a step of 2.5 kb was applied in the calculation of both the GC skew and the GC content.

The assembled genomes of the SOB (98.55% genome completeness; 11 scaffolds of 4.91 Mb; 5,518 genes) and the MOB (99.25% genome completeness; 10 scaffolds of 2.93 Mb; 3,102 genes) (**Table S4, Table S5, Table S6**) showed different genomic features in terms of GC content of the deviation from the average 37.22%, GC skew, and annotated Clusters of Orthologous Groups (COG) (**Fig. 2c**; for details of assembly and functional annotation see Supplementary Materials). Bacterial sequences were barely detectable in the scaffolds of *G. aegis*, suggesting a lack of horizontal gene transfer from the prokaryote. The CheckM analysis confirmed the high quality of the two symbionts genomes, as the potential contamination was low (3.25% in SOB, 1.67% in MOB, **Table S4**). In the metaproteomic analyses of the three parties, a total of 704 proteins were identified in the host (**Table S7**), 462 proteins in the SOB (**Table S8**), and 119 proteins (**Table S9**) in the MOB, which provide additional protein evidence to trace the metabolism of *G. aegis* holobiont.

### Evolution of Symbiosis in Deep-sea Peltospirid Snails

Molecular clock methods based on 1,066 single-copy genes suggests that the two deep-sea peltospirid snails, *G. aegis* and *C. squamiferum*, diverged approximately 117.94 million years ago (Ma; 95% confidence intervals: 68.73∼187.03 Ma) (**Fig. 3a**), which is consistent with previous evidence that the ancestor of peltospirids extends to the Late Cretaceous and the order Neomphalida likely had a mid-Jurassic origin^15^. Despite occupying the same microhabitat, the peltospirid snails *G. aegis* and *C. squamiferum* host different lineages of sulphur-oxidising endosymbionts (**Fig. 3b, Table S10**), consistent with the horizontal acquisition of symbionts^8,18^ and host-specific selectivity of symbionts relying on the pattern recognition receptors^22^. The host-specific selectivity for different lineages of gammaproteobacterial endosymbionts in peltospirids is in line with a previous study indicating that they have independent origins of endosymbioses^15^. In contrast, provannid *Alviniconcha* snails (*Alviniconcha strummeri, A. kojimai*) from vent fields at the Eastern Lau Spreading Center^8^ have been found to be capable of accepting a wide range of endosymbionts in each host species, and the endosymbiont has an impact on the holobiont’s niche utilisation^23^. This difference between peltospirids and provannids likely indicates that housing endosymbionts in an internal organ is associated with higher host selectivity of endosymbionts.

**Figure 3.**
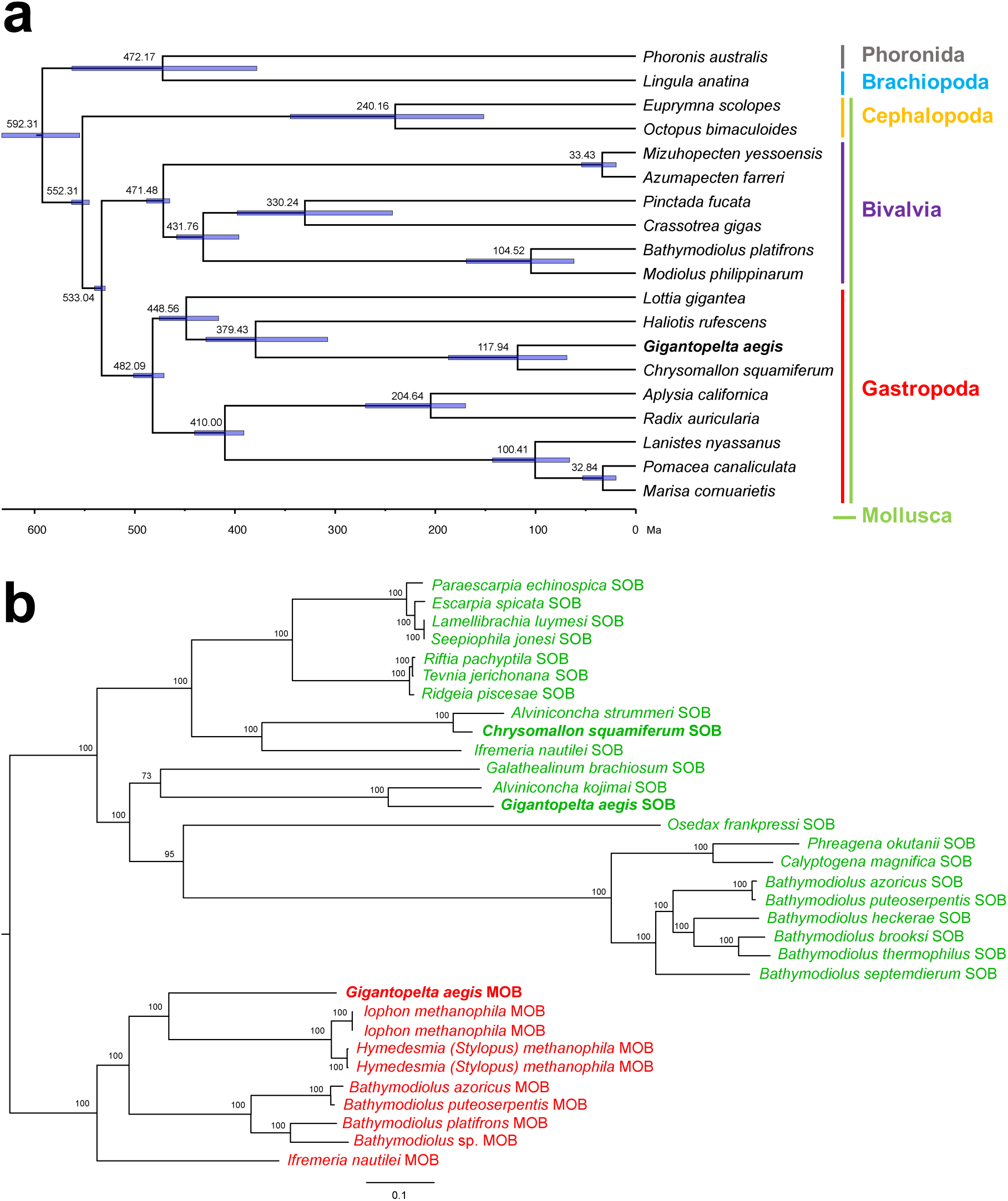
**a**. Time-calibrated phylogeny showing the estimated divergence time of 19 lophotrochozoan taxa, with a focus on 17 molluscs. *Phoronis australis* (Phoronida) and *Lingula anatina* (Brachiopoda) served as outgroups for the molluscs. A total of 1,066 single-copy orthologs were used. The purple horizontal lines indicate the 95% confidence intervals of the divergence times. For genome resources, fossil records and geographic events used in the time calibration see Supplementary Materials. **b**. Phylogenetic reconstruction of 32 symbionts in Gammaproteobacteria (See Supplementary Materials) from invertebrate taxa living in deep-sea chemosynthetic environments. A total of 424 single-copy orthologs were used to construct the tree. The phylogenetic tree includes two major lineages of the sulphur-oxidising bacteria (green) and the methane-oxidising bacteria (red).

Despite the divergent phylogenetic distances between the sulphur-oxidising endosymbionts in the two peltospirid snails, the gene content encoding for nutrition metabolism in the symbionts are similar, and the same goes for the two peltospirid hosts (**Table 1**). Divergence of hosts and their SOBs at the genome level together with the convergence of host and SOB gene content relating to nutrition again indicate that the peltospirid snails established their symbiotic relationships through convergent evolutionary pathways. Therefore, the selection of the symbionts may have been based on their capability of the biosynthesis of specific nutrients, especially key amino acids and vitamins that the hosts are unable to synthesise themselves.

**Table 1.**
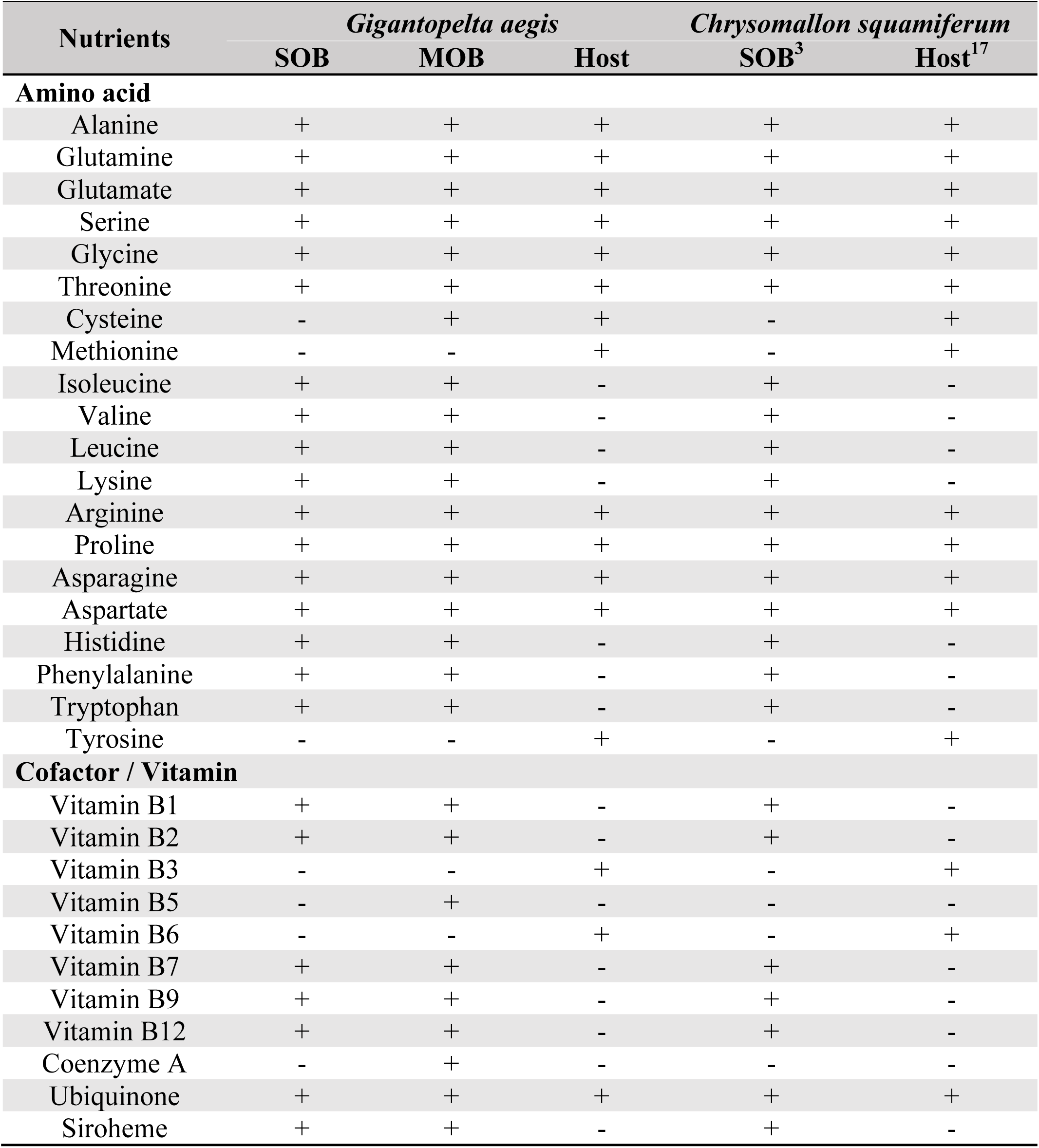
Genome capabilities of *Gigantopelta aegis* and *Chrysomallon squamiferum* holobionts^3,17^ in the biosynthesis of different nutrients, including amino acids, cofactors and vitamins (SOB: sulphur-oxidising endosymbiont, MOB: methane-oxidising endosymbiont).

| Nutrients | *Gigantopelta aegis* |  |  | *Chrysomallon squamiferum* |  |
| --- | --- | --- | --- | --- | --- |
|  | SOB | MOB | Host | SOB<sup>3</sup> | Host<sup>17</sup> |
| **Amino acid** |  |  |  |  |  |
| Alanine | + | + | + | + | + |
| Glutamine | + | + | + | + | + |
| Glutamate | + | + | + | + | + |
| Serine | + | + | + | + | + |
| Glycine | + | + | + | + | + |
| Threonine | + | + | + | + | + |
| Cysteine | - | + | + | - | + |
| Methionine | - | - | + | - | + |
| Isoleucine | + | + | - | + | - |
| Valine | + | + | - | + | - |
| Leucine | + | + | - | + | - |
| Lysine | + | + | - | + | - |
| Arginine | + | + | + | + | + |
| Proline | + | + | + | + | + |
| Asparagine | + | + | + | + | + |
| Aspartate | + | + | + | + | + |
| Histidine | + | + | - | + | - |
| Phenylalanine | + | + | - | + | - |
| Tryptophan | + | + | - | + | - |
| Tyrosine | - | - | + | - | + |
| **Cofactor / Vitamin** |  |  |  |  |  |
| Vitamin B1 | + | + | - | + | - |
| Vitamin B2 | + | + | - | + | - |
| Vitamin B3 | - | - | + | - | + |
| Vitamin B5 | - | + | - | - | - |
| Vitamin B6 | - | - | + | - | + |
| Vitamin B7 | + | + | - | + | - |
| Vitamin B9 | + | + | - | + | - |
| Vitamin B12 | + | + | - | + | - |
| Coenzyme A | - | + | - | - | - |
| Ubiquinone | + | + | + | + | + |
| Siroheme | + | + | - | + | - |

### Genetic Control of Symbiont Acquisition

The immune response of the host is critical for symbiont infection and colonisation. Among the shared gene families of *G. aegis* and 18 other lophotrochozoan genomes (**Fig. S8**), several immune-related gene families were expanded in *G. aegis* (**Table S11**). For example, the gene families fucolectin (FUCL) and galectin (Gal) were prominently expanded in *G. aegis* (**Fig. 4a**). These lectins allow host cells to bind with carbohydrate recognition domains on the bacterial surface, resulting in recognition of beneficial bacteria and elimination of detrimental microbes via antimicrobial activity^22^. In comparison to *C. squamiferum, G. aegis* harbours much more abundant pattern recognition receptors, including peptidoglycan recognition proteins (PGRP), toll-like receptors (TLR), fibrinogen-related proteins (FBG), and C-type lectins (CLEC) (**Fig. 5**). These receptors are thought to help other chemosymbiotic host animals, such as siboglinid tubeworms and bathymodiolin mussels, to respond to the microbes appropriately as well as assist the acquisition and maintenance of their symbiont populations^22^. Pattern recognition receptors that are highly expressed in the oesophageal glands are likely related to the recognition of the endosymbionts. From gene expression patterns, all PGRPs are highly expressed in the oesophageal glands of both peltospirid snails, suggesting PGRPs play an important role in the establishment of the symbiosis in deep-sea peltospirid snails. The abundance and high expression of PGRPs may help the host to regulate the symbiont population^22^ via the digestion process (**Table S12**), resulting in efficient utilisation of environmental chemical compounds whose concentrations are expected to fluctuate. Besides PGRPs, many other pattern recognition receptors, particularly those belonging to the gene family TLR, are highly expressed in the oesophageal gland of *G. aegis* compared to *C. squamiferum* (**Fig. 5**). Although *G. aegis* occupies the same habitat in the Longqi vent field as *C. squamiferum*, these additional pattern recognition receptors may enable *G. aegis* to host two distinct endosymbionts.

**Figure 4.**
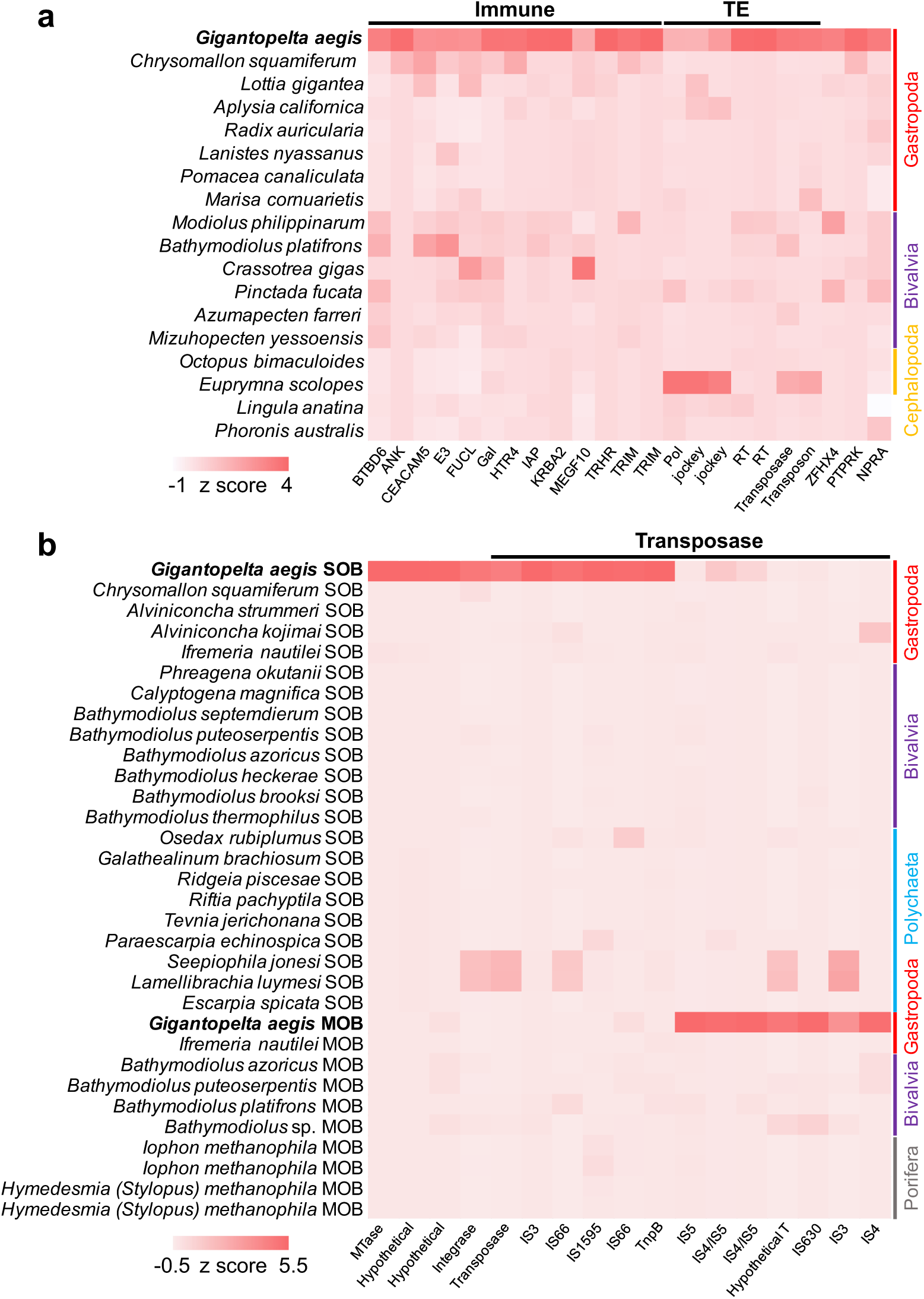
Heat maps showing the expanded gene families in *Gigantopelta aegis* and its two endosymbionts. **a**. Compared to other 15 molluscan taxa (systematic affinity at the class level shown on the right side), *Phoronis australis* (Phoronida), and *Lingula anatina* (Brachiopoda) with available genomes (used in the molecular clock analysis), the gene families expanded in *G. aegis* mainly include immune-related gene families (Immune) and transposable elements (TEs). **b**. Both the sulphur-oxidising endosymbionts and the methane-oxidising endosymbionts exhibit the transposases expansion. References are 30 symbionts in Gammaproteobacteria (See Supplementary Materials) from invertebrate taxa living in deep-sea chemosynthetic environments. The systematic affinity of the host taxa is shown on the right side.

**Figure 5.**
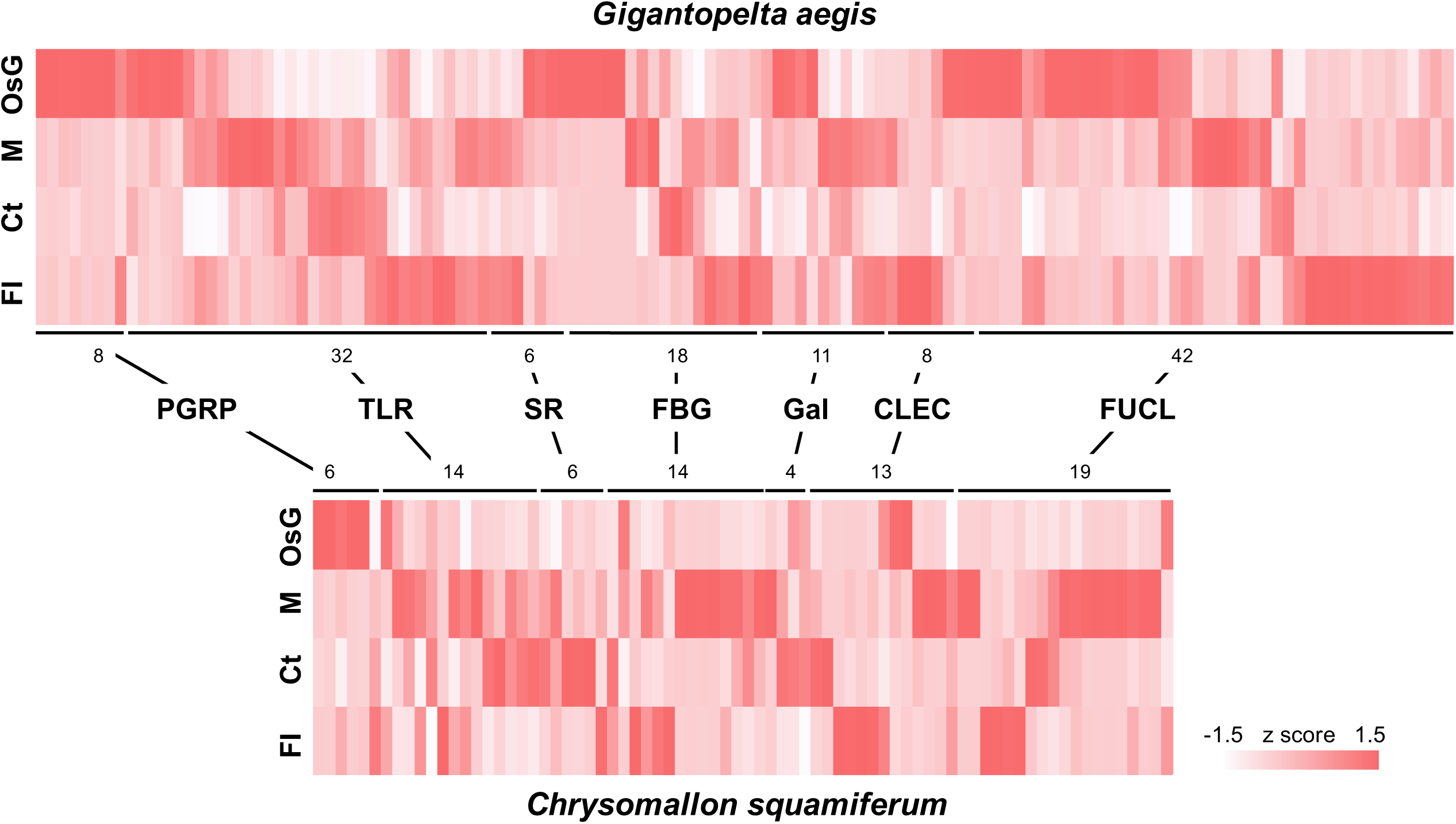
Two heat maps showing gene abundances and expression levels of pattern recognition receptors, including peptidoglycan recognition proteins (PGRP), toll-like receptors (TLR), scavenger receptor (SR), fibrinogen-related proteins (FBG), galectins (Gal), C-type lectins (CLEC), and fucolectin (FUCL), in *Gigantopelta aegis* (*n* = 4) and *Chrysomallon squamiferum* (*n* = 3)^17^ (OsG: oesophageal gland; M: mantle; Ct: ctenidium; FI: internal tissue of foot). The numbers near the lines indicate the gene numbers of the gene families.

There is also genomic evidence that the symbionts of the *G. aegis* peltospirids play a role in the invasion of the host. The peptidoglycan-associated lipoprotein Pal that can bind with TLR protein families^8^ was expressed in the transcriptomes and proteomes of the *G. aegis* SOB (**Table S5, Table S8**). Furthermore, both the SOB and the MOB of *G. aegis* contained OmpA family proteins and OmpH family proteins (**Table S5, Table S6**), which could interact with TLR and scavenger receptors to help the bacteria invade and adapt to the intracellular environment^24^. These symbiont attributes could help the host establish an endosymbiotic lifestyle.

### Energy Resources

Previous reports showed that the concentrations of reduced components (i.e., CH4, sulphur substances, and hydrogen) in the endmember fluid from the Longqi field are comparable or higher than those in the vents along the Central Indian Ridge^25,26,27^. Although methane is enriched in Longqi compared to other vents on the Central Indian Ridge^26^, further biogenic methane input from microbes living below and around the community may be sufficient to sustain the MOB. The blood vascular system of the host snail transports these essential dissolved compounds from the vent fluid, together with oxygen from the surrounding seawater, to the SOB and MOB **(Table S13)**. The dual symbionts are highly versatile in utilising those chemical substances, respectively (**Fig. 6a**). The central metabolism, associated gene expression levels, and protein abundance levels of the symbionts are summarised in **Figure 6**.

**Figure 6.**
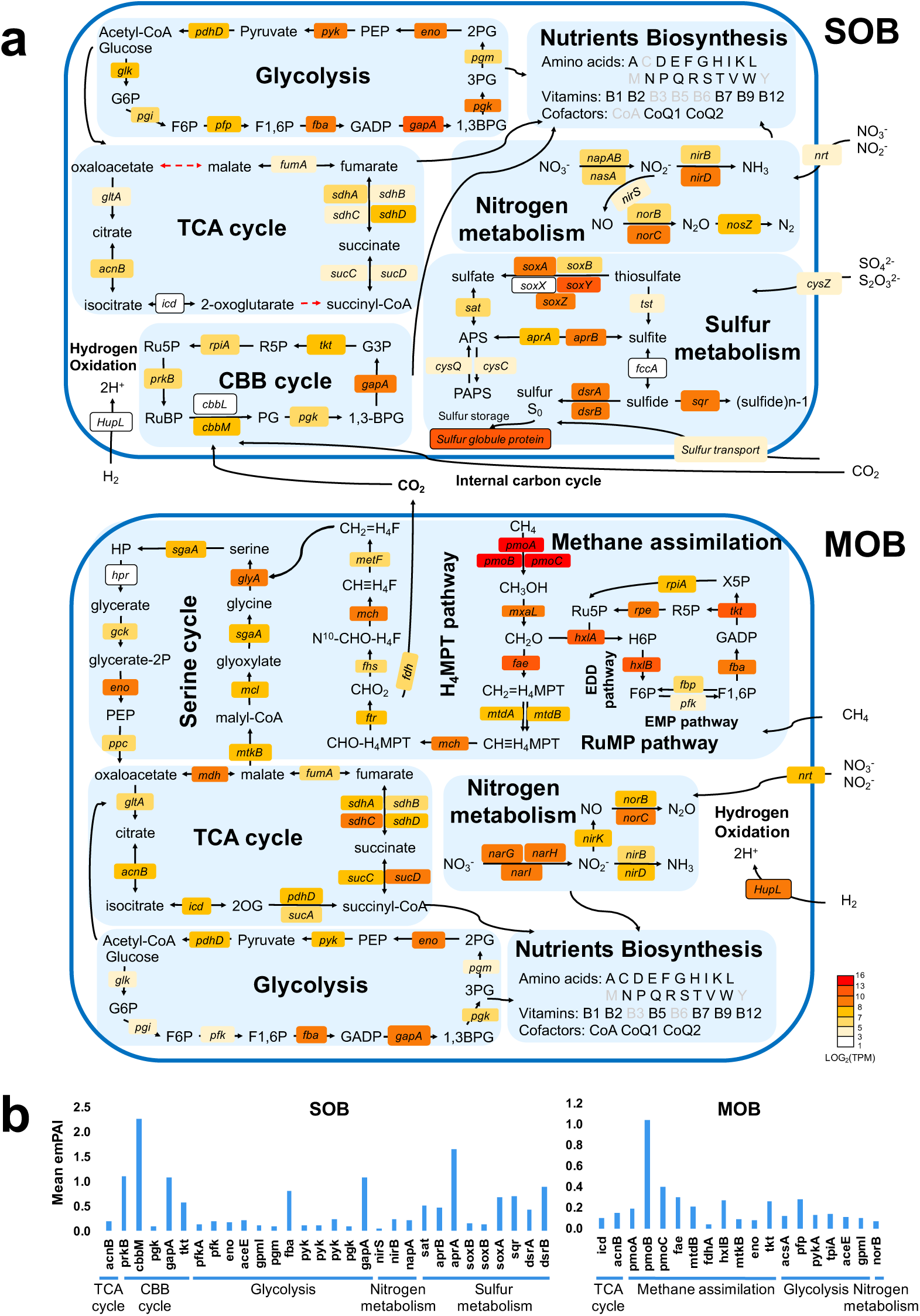
Central metabolism of the dual symbionts of *Gigantopelta aegis*. **a**. Central metabolism pathways in the sulphur-oxidising endosymbiont (SOB) and the methane-oxidising endosymbiont (MOB). The boxes represent genes involved in the respective process, and they are colour coded according to the Log-transformed normalized transcripts per kilobase million (TPM) value (*n* = 3) showing the gene expression level in the metatranscriptome analysis (colour from red to light white represents high expression level to low expression level; dash arrow represents that the gene is missing in the genome). **b**. Protein abundances involved in the central metabolism of the SOB (left) and the MOB (right). Protein abundances are reported as mean emPAI values (*n* = 3) assessed in the metaproteome analysis.

The genome data indicate that the sulphur metabolism in the SOB of *G. aegis* (**Fig. 6a**) is similar to that of the *C. squamiferum* snail SOB, *Alviniconcha* snail gamma SOB and siboglinid tubeworm gamma SOB^3,5^. The proteins of *soxA* (35th in proteome), *dsrB* (21th in proteome), *sqr* (34th in proteome), *aprA* (6th in proteome), *aprB* (63th in proteome), and *sat* (57th in proteome) (**Fig. 6**) have a high expression level, indicating their active engagement in oxidising thiosulfate, sulphite, and sulphide in the environment and to detoxify sulphide for the holobiont.

Like many sulphur-oxidising endosymbionts of vent molluscs^3,4,28^, the SOB of *G. aegis* relies on the complete Calvin-Benson-Bassham (CBB) cycle to fix CO2 by using a form II ribulose-bisphosphate carboxylase (the protein of *cbbM* gene: 2nd in proteome) that was found the highest protein expression level and lacks a complete rTCA cycle (**Fig. 6**). Phosphoribulokinase (the protein of *prkB* gene: 15th in proteome), type I glyceraldehyde-3-phosphate dehydrogenase (the protein of *gapA* gene: 16th in proteome), and transketolase (the protein of *tkt* gene: 43th in proteome) playing important roles in CBB cycle for carbon fixation also exhibited high protein expression levels in the SOB (**Fig. 6**).

The MOB of *G. aegis* has neither a complete CBB cycle nor an rTCA cycle pathway for CO2 fixation. However, the *G. aegis* MOB possesses the methane monooxygenase operon (*pmoA* gene: 5th in transcriptome, 50th in proteome; *pmoB* gene: 6th in transcriptome, 1st in proteome; *pmoC* gene: 1st in transcriptome, 10th in transcriptome) with top 1% gene expression enabling it to oxidise methane (**Fig. 6b, Fig. S9**), confirming that methane oxidation is among the most prominent metabolic processes in the MOB. In the tetrahydromethanopterin (H4MPT) pathway, the high expression level of the formaldehyde activating enzyme (*fae* gene: 65th in transcriptome, 20th in proteome) indicates an active generation of CO2 via formaldehyde oxidation. Then CO2 could be redirected to the CBB cycle of the SOB to form an internal carbon cycle among the symbionts. Internal carbon cycling among symbionts has been detected in the internal symbiotic organs (trophosomes) of vent siboglinid tubeworms^29^. This would allow the symbionts to maximise carbon efficiency within the trophosome-like oesophageal glands and trophosomes in the absence of direct interaction with CO2-rich seawater and vent fluids.

Aerobic methanotrophs are classified into two groups based on their formaldehyde assimilation pathways: Type I group (part of Gammaproteobacteria) using a ribulose monophosphate (RuMP) cycle and Type II group (part of Alphaproteobacteria) using a serine cycle. Methane-oxidising symbionts usually belong to Type I and only use the RuMP cycle; examples include the methane-oxidising symbionts of deep-sea *Bathymodiolus* mussels, snails, and sponges (referred-to genomes in **Fig. 3b**). The MOB of *G. aegis*, however, surprisingly possesses a complete serine cycle in addition to a complete RuMP cycle (**Fig. 6a)**. This can mitigate the accumulation and toxicity of formaldehyde when formaldehyde is in excess for the RuMP pathway. In the RuMP cycle, high expression of 3-hexulose-6-phosphate synthase (*hxlA* gene: 20th in transcriptome) and 6-phospho-3-hexuloisomerase (*hxlB* gene: 25th in proteome) in the Entner-Doudoroff (EDD) pathway indicate an important role for formaldehyde oxidation (**Fig. 6)**. Furthermore, the MOB genome possesses an additional Embden-Meyerhof-Parnas (EMP) pathway for glycolysis (**Fig. 6a)**, absent in other methane-oxidising endosymbionts (referred in **Fig. 3b**). These results indicate that MOB of *G. aegis* are highly efficient and versatile in assimilating carbon sources from methane.

### Respiration

Both the SOB and the MOB housed in the oesophageal glands of *G. aegis* are capable of using nitrate as an electron acceptor for anaerobic respiration. Their genomes encode genes for nitrate reduction, such as nitrate reductase (*napAB* and *nasA* in SOB, *narGHI* in MOB), nitrite reductase (*nirBDS* in SOB, *nirBDK* in MOB), and nitric oxide reductase (*norBC* in the SOB and the MOB) (**Fig. 6, Fig. S10**), which are also found in sulphur-oxidising symbionts of tubeworms, mussels, and other gastropods but not in the methane-oxidising symbionts of mussels and sponges^3,5,28,30^. As the *G. aegis* symbionts rely on the host’s blood vessels for oxygen supply while input from the hydrothermal vent fluctuates due to the constant interplay between the oxygen-poor vent fluid and the oxygen-rich seawater, there may be periods when the oxygen supply runs low in the oesophageal gland. Indeed, hemocyanin, hemoglobin, and globin-like genes playing a role in oxygen binding and transport have lower gene expression in the oesophageal gland and higher expression in the foot and mantle while five sialin genes used for transporting nitrate were highly expressed in the oesophageal gland (**Fig. S11**). Therefore, these symbionts might have developed abilities to avoid oxygen competition with the host, especially under potential periodic hypoxic conditions.

### Holobiont Nutritional Interdependence

The SOB do not have *mdh, pdhD* and *sucA* genes in the TCA cycle while the homologous genes of those missing in the SOB are highly expressed in the MOB. The SOB does have a C4-dicarboxylate TRAP transport system (DstPQM) that would allow the uptake of four-carbon compounds involved in the TCA cycle^8^, indicating that the SOB likely replenishes the missing intermediates of the TCA cycle from the other symbiont partner.

Both the *G. aegis* and *C. squamiferum* host genomes lack the capability of the biosynthesis of 7 amino acids (isoleucine, valine, leucine, lysine, histidine, phenylalanine, and tryptophan), 6 vitamins (vitamin B1, vitamin B2, vitamin B5, vitamin B7, vitamin B9, and vitamin B12), and 2 cofactors (siroheme and coenzyme A; **Table 1**). However, these nutrients, except vitamin B5 and coenzyme A, can be obtained from their SOB through host lysis symbiont cells (**Table S12**). The MOB of *G. aegis* has a complete pathway to synthesise vitamin B5 and coenzyme A, whereas the SOBs of two peltospirids lacks a key gene 2-dehydropantoate 2-reductase (*panE*) for generating pantoate, the key substrate for producing pantothenate (**Fig. S12**). We speculate two possibilities: 1) the SOB, particularly of *C. squamiferum*, has a replacement of 2-dehydropantoate 2-reductase or replenishment for the biosynthesis of pantothenate and coenzyme A like in the pea amphid holobiont^31^, or 2) in the *G. aegis* holobiont, MOB serves as the major source for providing pantothenate and coenzyme A to both the SOB and the host. In return, the host can provide the SOB and the MOB with methionine, tyrosine, vitamin B3, and vitamin B6 through the symbionts’ ATP-binding cassette transporters of amino acids. These host-supplied nutrients are critical for fulfilling the nutritional demands of the endosymbionts housed in an internal organ lacking direct contact with the outside world.

### Transposase Expansion in Three Partners of the *Gigantopelta aegis* Holobiont

The enrichment of the transposable elements (TE) provides beneficial genomic plasticity for adaptation to an intracellular symbiotic life by regulating nearby genes^32^. In the results from the gene family analysis, expansion of TEs were found in all three parties comprising *G. aegis* holobiont (**Fig. 4**). For the host *G. aegis*, these expansions correspond to 342 transposase genes (1.6% of total host genes) and 510 DNA transposons (2.4% of total host genes); for SOB symbionts, these expansions correspond to 704 transposase genes (12.8% of total SOB genes); and for the MOB symbionts, these expansions correspond to 345 transposase genes (11.1% of total MOB genes). Numerous transposase genes were unusually inserted into the flagellar operons and chemotaxis operons playing an important role in host infection by both SOB and MOB of *G. aegis*. This may contribute to the regulation of the symbionts motility for infection and colonisation in the host environment. To our knowledge, the expansion of transposase genes occurring across multiple symbiotic partners within the same holobiont has not been reported in any marine invertebrate holobionts.

## Conclusions

We revealed that the deep-sea peltospirid snail *Gigantopelta aegis* endemic to hydrothermal vent exhibits a dual symbiotic lifestyle, which includes dominant sulphur-oxidising Gammaproteobacteria and less dominant methane-oxidising Gammaproteobacteria (Type I methanotroph). Furthermore, we provide a high-quality hologenome including a chromosome-level assembly for the snail host and an in-depth analysis of the genomic interdependencies among the trinity of symbiotic partners in this holobiont, finding an intimate mutualistic relationship with complementarity in nutrition and metabolic codependency. The two endosymbionts live in an organ lacking direct interaction with the ambient environment. Both symbionts as well as the host are highly versatile with regards to the transportation and utilisation of chemical energy, increasing the efficiency of carbon fixation by forming an internal carbon cycle among the two symbionts. Although *G. aegis* and another chemosymbiotic peltospirid snail *Chrysomallon squamiferum* occupy the same habitat in the Longqi vent field, *C. squamiferum* has a single sulphur-oxidising symbiont. The fact that the sulphur-oxidising endosymbionts of both snails are not particularly closely related phylogenetically but have essentially the same capabilities of the biosynthesis of specific nutrition suggest that this may be a key criterion for the selection of symbionts by the snail hosts.

## Materials and Methods

### Deep-Sea Sampling

*Gigantopelta aegis* snails were collected by the manned submersible HOV *Jiaolong* on-board the R/V *Xiangyanghong 9* cruise 35II from Longqi hydrothermal vent field (37.7839°S, 49.6502°E; 2,761 m depth) on the Southwest Indian Ridge in January 2015. Snails were immediately flash-frozen in liquid nitrogen once they were recovered on the ship. They were then transferred to −80 °C until DNA and RNA extraction. Specimens of *G. aegis* for Hi-C sequencing and fluorescence *in situ* hybridisation (FISH) experiments were collected also from Longqi (‘Tiamat’ chimney) during Leg 3 of the COMRA R/V *Dayang Yihao* expedition 52 in April 2019. Tissue from the foot was immediately dissected from one individual, cut up, washed in phosphate-buffered saline buffer, and stored at −80 °C until Hi-C library preparation. The oesophageal gland tissue was dissected from the same individual and fixed in 4% paraformaldehyde overnight, followed by dehydration through an ethanol series (20%, 40%, 60%, and 80%) for 15 minutes each and stored at −80 °C until fluorescence *in situ* hybridisation experiments. Samples for transmission electron microscopy (TEM) were immediately fixed in 10% buffered formalin after recovery on-board the research vessel. Identity of the specimens was confirmed by both mitochondrial *COI* sequences and morphology.

### TEM

The oesophageal gland tissue was dehydrated using a series of graded acetone and then transferred into Epon resin (Sigma-Aldrich) for embedding. An ultramicrotome (Reichert Ultracut S, Leica) was used to slice ultrathin (70 nm) sections that were used for staining in 2% aqueous uranyl acetate with lead stain solution (0.3% lead acetate and 0.3% lead nitrate, Sigma-Aldrich). The presence of intracellular symbionts was confirmed using a Tecnai 20 Transmission Electron Microscopy (FEI) at an acceleration voltage of 120 kV. For more detailed methods for thin-section preparation see ref 15.

### FISH

The oesophageal gland tissues were dehydrated in 100% ethanol and embedded in Shandon Cryomatrix Frozen Embedding Medium (6769006; Thermo Fisher Scientific). After full embedding, they were quickly frozen in dry ice. Cryostat sections 10 µm in thickness (CryoStar NX70; Thermo Fisher Scientific) were cut and mounted on glass slides. Two 16S rRNA-based probes were designed to target the endosymbionts specifically. The Cy3-labelled SOB1 probe (5’-AGCATATTAAACTTGTACCC-3’) was used to target the sulphur-oxidising endosymbiont, and the Cy5-labelled MOB1 probe (5’-CGTGTGTTTTCCTCCCTTCT-3’) was used to bind the methane-oxidising endosymbiont. The sections were rehydrated in a decreasing ethanol series (95%, 80%, and 70%) for 15 minutes each and hybridised at 46 °C with 50 ng/ml of each probe in a hybridisation buffer (0.9 M NaCl, 0.02M Tris-HCl, 0.01% sodium dodecyl sulphate and 20% formamide) for 3 hours. Then, the slides were washed in a wash buffer (0.1 M NaCl, 0.02 M Tris-HCl, 0.01% sodium dodecyl sulphate, 5 mM EDTA) at 48 °C for 15 min. Two drops of 4′,6-diamidino-2-phenylindole (DAPI) were added on each slide followed by incubation at room temperature for 3 min. After washing and air drying, the slides were mounted by coverslips with SlowFade™ Diamond Antifade Mountant medium (Invitrogen, Carlsbad, CA, USA). Images were taken using a confocal microscope (Leica Microsystems, Wetzlar, Germany) with a 3D model, and post-processed by LAS X software (Leica Microsystems, Wetzlar, Germany).

### DNA Extraction and Illumina Sequencing of the Hologenome

The MagAttract High-Molecular-Weight DNA Kit (QIAGEN, Hilden, Netherlands) was used to extract high-molecular-weight genomic DNA from the oesophageal gland and the foot separately for sequencing of the genome of the symbionts and the host, respectively, following manufacturer’s protocols. The extracted DNA was further purified by Genomic DNA Clean & Concentrator™-10 kit (ZYMO Research, Irvine, CA, USA). The DNA was finally eluted in 10mM Tris-HCl buffer (pH=8.5). The quality of the genomic DNA was evaluated using a BioDrop µLITE (BioDrop, Cambridge, UK) with the OD 260/280 of 1.8 and the OD 260/230 of 2.0–2.2. The DNA concentration was assessed using a Quibit™ 3 Fluorometer (Thermo Fisher Scientific, Singapore). The sizes of DNA fragments were assessed by pulsed-field gel electrophoresis. Approximate 1 µg DNA of each tissue, including foot and oesophageal gland, were used for the short-insert library (350 bp and 500 bp) in an Illumina NovaSeq 6000 platform to generate 171 Gb and 49 Gb paired-end reads with a length of 150 bp, respectively.

### Oxford Nanopore Technologies (ONT) library preparation and MinION sequencing

The DNA of the oesophageal gland was used to prepare the long-read library (Oxford Nanopore MinION, UK) for genome sequencing of endosymbionts. To construct the MinION library, a NEBNext Ultra II End-Repair/dA-tailing Module (NEBE7546) kit was used to perform DNA repair and end-prep. A Ligation Sequencing Kit (SQK-LSK109) was used to perform adaptor ligation and clean-up with the large DNA fragments. A Flow Cell Priming Kit (EXP-FLP001) was used to prepare the priming buffer to prime the MinION flow cells (FLO-MIN106D R9 version). The Library Loading Bead Kit R9 version (EXP-LLB001) was used to assist in loading the libraries into the flow cells. Sequencing was started immediately in the MinION™ portable single-molecule nanopore sequencer (Oxford Nanopore Technologies Limited, UK) using the software MinKNOW version 3.1.8 (Oxford Nanopore Technologies Limited, UK). The read event data was base-called by Oxford Nanopore basecaller Guppy version 2.1.3^33^ with default settings.

### Single-Molecule Real-Time (SMRT) Library Construction and PacBio Sequel Sequencing

For genome sequencing of the host animal, the purified DNA of the foot from one individual was utilised to construct PacBio single-molecule real-time (SMRT) library (Pacific Biosciences, USA). BluePipin (Sage Science, USA) was utilised to select long DNA fragments with sizes between 8 kb to 50 kb for PacBio library construction. Universal hairpin adapters were used to ligate the DNA fragments. The adapter dimers and hairpin dimers were both removed using magnetic bead in PacBio’s MagBead kit. Exonucleases were further used to remove the failed ligation DNA fragments. The purified library was sequenced by binding the sequencing polymerase in the SMRT sequencing cells and generated long subreads with a length distribution (**Fig. S13)**.

### Hi-C Sequencing

To construct Hi-C library, the frozen foot tissue of *G. aegis* was thawed slowly on ice and suspended in 45 mL of 37% formaldehyde in serum-free Dulbecco’s modified Eagle’s medium (DMEM) for chromatin cross-linking. After incubation at room temperature for 5 minutes, glycine was added to quench the formaldehyde, followed by incubation at room temperature for another 5 minutes and subsequently on ice for over 15 minutes. The cells were further lysed in pre-chilled lysis buffer (10 mM NaCl, 0.2% IGEPAL CA-630, 10 mM Tris-HCl, and 1× protease inhibitor solution) using a Dounce homogenizer. The chromatin was digested by Mbo I restriction enzyme, marked with biotin and ligated^34^. The Hi-C library was sequenced on a NovaSeq 6000 platform (Illumina, USA) and generated reads with a length of 150 bp. The genome sequencing strategy of the host was summarised in **Table S14**.

### RNA Extraction, Metatranscriptome Sequencing, and Eukaryotic Transcriptome Sequencing

The total RNA of oesophageal glands dissected from three individuals of *Gigantopelta aegis* were separately extracted using the TRIzol reagent (Invitrogen, Carlsbad, CA, USA). The quality and quantity of RNA were checked by agarose gel electrophoresis and BioDrop µLITE (BioDrop, Cambridge, UK), respectively. To enrich the mRNA of endosymbionts, the rRNA of both eukaryote and bacteria were removed by both the Ribo-Zero™ Magnetic Kit (Human/Mouse/Rat) (Epicenter, USA) and the Ribo-Zero™ Magnetic Kit (Bacteria) (Epicenter, USA) following the manufacturers’ protocols. The remaining RNA was used to construct cDNA libraries for metatranscriptome sequencing. The sequencing data are shown in **Table S15**.

Four *G. aegis* individuals were dissected to obtain different organs (foot, mantle, ctenidium, oesophageal gland, etc) for eukaryotic transcriptome sequencing (**Table S15**). Total RNA of these samples were separately extracted using TRIzol reagent (Invitrogen, Carlsbad, CA, USA) following the manufacturers’ protocol. mRNA from each host tissue was enriched by Oligo-dT probes and further transferred to cDNA for eukaryotic library construction. All cDNA libraries were sequenced in a NovaSeq 6000 Illumina platform to generate paired-end reads with a length of 150 bp.

### Hologenome Assembly and Scaffolding

Adaptors and low-quality bases of raw Illumina sequencing reads were trimmed using Trimmomatic version 0.36^35^ with the settings “ILLUMINACLIP: Truseq3-PE-2.fa”. To obtain the genome of symbionts, clean Illumina reads from the oesophageal gland were used for initial assembly in metaSPAdes^36^ with a series of kmers of 51, 61, 71, 81, and 91 bp. Illumina short reads belonging to the symbionts were grouped by genome binning according to both the sequencing coverage and the GC content of the initial contigs^5^. The grouped Illumina short reads and all ONT sequencing reads were reassembled together by Unicycler version 0.4.7^37^. The assembled contigs were grouped again via genome binning. Each group of contigs were further used in genome scaffolding by SSPACE-LongRead version 1.1^38^ with the ONT reads and mapped PacBio reads that are over 10 kb. CheckM version 1.0.13^39^ was used to assess the completeness and the contamination of the assembled bacterial genomes.

Several assembly pipelines were used for assembly, and the best contig assembly result was generated by “Canu correction + wtdbg2” (**Table S16**). PacBio subreads over 4 kb were corrected by Canu version 1.7.1^40^ with settings of “genomeSize = 1.27 Gb, corMhapSensitivity = normal, corMinCoverage = 0, corMaxEvidenceErate = 0.15, correctedErrorRate = 0.065, minReadLength = 8000” and then used in genome assembly using wtdbg2 version 2.1^41^ with settings of “-e 2 --tidy-reads 5000 -S 1 -k 15 -p 0 --rescue-low-cov-edges --aln-noskip”. To improve the accuracy of the initially assembled contigs, two polishing rounds using Racon version 1.3.1^42^ were performed and followed by an additional round using Pilon version 1.22^43^ with Illumina short reads that were mapped by Bowtie2 version 2.3.5^44^ in the “--sensitive” mode. Bacterial contamination was further filtered using MaxBin version 2.2.5^45^.

Raw Hi-C sequencing reads were first trimmed by Trimmomatic version 0.36^35^. To remove the invalid pairs of Hi-C reads without effective ligation, the reads were mapped to the corrected assembled contigs using Bowtie2 version 2.3.5^44^, and Hi-C contact maps were generated based on the mapped reads using HiC-Pro^46^. Juicer version 1.5^47^ was used to filter and deduplicate. The remaining valid reads were used for contig scaffolding using the 3D *de novo* assembly (3D-DNA) pipeline^48^ in diploid mode. The scaffolds were manually corrected based on the Hi-C contact maps (**Fig. S14**). The continuity and completeness of the assembled genome were evaluated with assemblathon_stats.pl (https://github.com/ucdavis-bioinformatics/assemblathon2-analysis/blob/master/assemblathon_stats.pl) and BUSCO version 3.0.2^49^ using the “metazoa_odb9” database, respectively.

### Genes Prediction and Functional Annotation

The coding sequences and proteins of symbionts genomes were predicted by Prodigal version 2.6.3^50^ with default settings. The host genome was first hard-masked in the repetitive regions and then trained with the boundary of the introns and exons in order to predict coding genes of *G. aegis* (for details see Supplementary Materials).

Repeats in the *Gigantopelta aegis* genome were *de novo* identified and classified using RepeatModeler version 1.0.11 (http://www.repeatmasker.org/RepeatModeler/) pipeline and soft-masked using RepeatMasker version 4.0.8 (http://www.repeatmasker.org/RMDownload.html) with the parameter “-xsmall” (for details see Supplementary Materials).

To predict the gene models, we run two rounds of the MAKER pipeline version 2.31.10^51^ on the soft-masked genome. To guide the prediction, we used *de novo* assembled transcripts, Metazoan protein sequences downloaded from the Swiss-Prot database, as well as protein sequences of *Chrysomallon squamiferum*^17^ (for details see the Supplementary Materials).

The predicted protein sequences were searched against NCBI Non-Redundant (NR) database using BLASTp with an *E*-value cutoff of 1e-5. The sequences were also used to search the Kyoto Encyclopedia of Genes and Genomes (KEGG) database via KEGG Automatic Annotation Server (KAAS) using bi-directional best hit methods of BLASTp^52^. The protein sequences of the host were further searched against the EuKaryotic Orthologous Groups (KOG) database. The protein sequences of symbionts were annotated against the functional Clusters of Orthologous Groups (COG) using eggNOG-mapper v2^53^. Pfam database^54^ with hmmscan (http://hmmer.org/) was used to identify the functional domains of proteins.

### Gene Family and Phylogenomic analysis

To perform gene family analysis of *G. aegis*, Orthofinder version 2.3.3^55^ was used to identify the orthologous groups shared between proteins of *G. aegis* and those of other molluscan genomes (see Supplementary Materials). Only single-copy orthologs were used to construct the phylogenetic tree. The protein sequences of each gene were aligned separately using MAFFT v7.407^56^ and were concatenated for phylogenetic analysis using FastTree version 2.1.10^57^ with partitions. The software MCMCtree^58^ was used to yield the time-calibrated tree by calibrating the phylogenetic tree with seven fossil records and geographic events (see Supplementary Materials).

Both CAFE 3^59^ and Fisher’s Exact Test were used for gene family analysis. The time-calibrated tree and gene family numbers shared by these species were used for gene family analysis in CAFE 3^59^. The Fisher’s Exact Test was also used for gene family analysis based on the gene family numbers of *G. aegis* and the average gene family numbers of other species included. Only the gene families with a FDR corrected *P-*value smaller than 0.05 were considered as expanded or contracted.

To explore the phylogenetic relationship of the symbionts, 30 genomes of bacterial symbionts belonging to Gammaproteobacteria from deep-sea invertebrate taxa were included (details of referred symbionts see **Table S17**). Same pipelines of orthologs cluster and sequence alignments of *G. aegis* were applied in the symbionts analyses. The IQ-TREE multicore version 1.6.10^60^ with LG+I+G4+F were used to perform the phylogeny analysis of the symbionts with 1000 ultrafast bootstraps.

### Gene Expression Analysis of the Holobiont

The coding DNA sequences of the predicted genes were used as references to assess gene expression levels. Kallisto version 0.45.1^61^ was used to calculate transcripts per million (TPM) value according to the mapped reads count for each gene. In symbionts, the TPM values were directly used for gene expression level comparison. For the host, the highly expressed genes in the oesophageal gland were identified by differential expression analysis versus foot, ctenidium, and mantle (*n* = 4) in edgeR^62^. If genes had a fold change larger than two and a significant FDR *P*-value (< 0.05), they were designated as highly expressed genes. To compare the genes expression of *G. aegis* and *C. squamiferum*, the same analysis pipeline was applied to the transcriptome sequencing data of foot tissue, ctenidium, mantle, and oesophageal gland from the *C. squamiferum*^17^.

### Synteny Analysis

Proteins of *G. aegis* and *C. squamiferum* were searched against each other via BLASTp with an *E*-value cutoff of 1e-5. The hits were used to detect collinear blocks and synteny shared between these two deep-sea gastropods by MCScanX^63^. JCVI (https://github.com/tanghaibao/jcvi) was utilised to generate the chromosome-scale synteny plot.

### Metaproteomics

The oesophageal glands from three *Gigantopelta aegis* individuals were used for protein extraction using the methanol-chloroform method^64^. SDS-PAGE gel was used to separate different size (ranging from 10 kDa to 150 kDa) of ∼30 µg extracted protein from each sample and stained by colloidal coomassie blue. The peptide for LC-MS/MS was obtained through protein reduction, alkylation and digestion, peptide extraction and further dry. Dionex UltiMate 3000 RSLCnano coupled with an Orbitrap Fusion Lumos Mass Spectrometer (Thermo Fisher) was utilised to analyse each protein fraction. The search database contains the protein sequences predicted from the genome and the corresponding reversed sequences (decoy) of both *Gigantopelta aegis* and its two endosymbionts. Mascot version 2.3.0 was used to identify and quantify the protein via the raw mass spectrometry data. Proteins were identified with the assigned peptides’ identification confidence level over 0.95 and a false discovery rate of less than 2.5% (see Supplementary Materials).

## Acknowledgements

We thank captain and crew of the R/V *Xiangyanghong 9* and pilots of HOV *Jiaolong* for their great support during the research cruise DY35th-II, and captain and crew of the R/V *Dayang Yihao* as well as the operation team of the ROV *Sea Dragon* III during the third leg of the China Ocean Mineral Resources Research and Development Association DY52nd cruise. Dr. Jack C.H. Ip and Dr. Ting Xu from the Hong Kong Baptist University are gratefully acknowledged for their helpful comments. This work was supported by grants from China Ocean Mineral Resources Research and Development Association (DY135-E2-1-03), the Hong Kong Branch of Southern Marine Science and Engineering Guangdong Laboratory (Guangzhou) (SMSEGL20SC01), Southern Marine Science and Engineering Guangdong Laboratory (Guangzhou) (GML2019ZD0409), and Major Project of Basic and Applied Basic Research of Guangdong Province (2019B030302004-04) awarded to P-YQ.

## Data Availability

All sequencing data, assembly data, predicted genes, and proteins of *Gigantopelta aegis* and its two symbionts were submitted to the database of National Centre for Biotechnology Information under BioProject PRJNA612619.

## Author Contributions

P-YQ conceived the project. YL, JS, CC, and P-YQ designed the experiments. YZ, JS, and YS collected the samples of *Gigantopelta aegis* gastropods. CC, JS, YL, and YY dissected the *Gigantopelta aegis* samples. CC performed TEM experiments. YL, JS, AC, and WZ contributed to the FISH experiments. YL, SB, RL, and JS performed host genome assembly. YL and YS performed the gene prediction of *G. aegis*. RL drew Circos plots of the hologenome. YL and JS contributed to phylogenomic analysis and calibration of the molecular clock of the host. YL, KZ, YY, and WZ contributed to the symbiont genome assembly. WCW, YHK, JS, and YL performed the proteome analysis. YL performed the DNA extraction, RNA extraction, ONT sequencing, gene expression analysis, gene family analysis, synteny analysis, immune response analysis, and drafted the manuscript. J-WQ and CVD contributed to manuscript editing. All authors contributed to the manuscript and approved it for submission and publication.

## References

1. Robbins, S. J. et al. A genomic view of the reef-building coral *Porites lutea* and its microbial symbionts. Nat. Microbiol. 4, 2090–2100 (2019).

2. Dubilier, N., Bergin, C. & Lott, C. Symbiotic diversity in marine animals: the art of harnessing chemosynthesis. Nat. Rev. Microbiol. 6, 725–740 (2008).

3. Nakagawa, S. et al. Allying with armored snails: the complete genome of gammaproteobacterial endosymbiont. ISME J. 8, 40–51 (2014).

4. Newton, I. L. G. et al. The *Calyptogena magnifica* chemoautotrophic symbiont genome. Science 315, 998–1000 (2007).

5. Yang, Y. et al. Genomic, transcriptomic, and proteomic insights into the symbiosis of deep-sea tubeworm holobionts. ISME J. 14, 135–150 (2019).

6. Lan, Y. et al. Host-symbiont interactions in deep-sea chemosymbiotic vesicomyid clams: insights from transcriptome sequencing. Front. Mar. Sci. 6, 680 (2019).

7. Fujiwara, Y., Kato, C., Masui, N., Fujikura, K. & Kojima, S. Dual symbiosis in the cold-seep thyasirid clam *Maorithyas hadalis* from the hadal zone in the Japan Trench, western Pacific. Mar. Ecol. Prog. Ser. 214, 151–159 (2001).

8. Beinart, R. A., Luo, C., Konstantinidis, K., Stewart, F. & Girguis, P. R. The bacterial symbionts of closely related hydrothermal vent snails with distinct geochemical habitats show broad similarity in chemoautotrophic gene content. Front. Microbiol. 10, 1818 (2019).

9. Ansorge, R. et al. Functional diversity enables multiple symbiont strains to coexist in deep-sea mussels. Nat. Microbiol. 4, 2487–2497 (2019).

10. Geier, B. et al. Spatial metabolomics of in situ host–microbe interactions at the micrometre scale. Nat. Microbiol. 5, 498–510 (2020).

11. Sun, J. et al. Adaptation to deep-sea chemosynthetic environments as revealed by mussel genomes. Nat. Ecol. Evol. 1, 121 (2017).

12. Won, Y. J. et al. Environmental acquisition of thiotrophic endosymbionts by deep-sea mussels of the genus *Bathymodiolus*. Appl. Environ. Microbiol. 69, 6785–6792 (2003).

13. Goffredi, S. K., Warén, A., Orphan, V. J., Van Dover, C. L. & Vrijenhoek, R. C. Novel forms of structural integration between microbes and a hydrothermal vent gastropod from the Indian Ocean. Appl. Environ. Microbiol. 70, 3082–3090 (2004).

14. Chen, C., Linse, K., Copley, J. T. & Rogers, A. D. The ‘scaly-foot gastropod’: a new genus and species of hydrothermal vent-endemic gastropod (Neomphalina: Peltospiridae) from the Indian Ocean. J. Molluscan Stud. 81, 322–334 (2015).

15. Chen, C., Uematsu, K., Linse, K. & Sigwart, J. D. By more ways than one: rapid convergence at hydrothermal vents shown by 3D anatomical reconstruction of *Gigantopelta* (Mollusca: Neomphalina). BMC Evol. Biol. 17, 62 (2017).

16. Chen, C., Linse, K., Uematsu, K. & Sigwart, J. D. Cryptic niche switching in a chemosymbiotic gastropod. Proc. Biol. Sci. 285, 20181099 (2018).

17. Sun, J. et al. The scaly-foot snail genome and the ancient origins of biomineralised armour. Nat. Commun. 11, 1657 (2020).

18. Heywood, J. L., Chen, C., Pearce, D. A. & Linse, K. Bacterial communities associated with the Southern Ocean vent gastropod, *Gigantopelta chessoia*: indication of horizontal symbiont transfer. Polar Biol. 40, 2335–2342 (2017).

19. Sun, J. et al. Signatures of divergence, invasiveness, and terrestrialization revealed by four apple snail genomes. Mol. Biol. Evol. 36, 1507–1520 (2019).

20. Salerno, J. L. et al. Characterization of symbiont populations in life-history stages of mussels from chemosynthetic environments. Biol. Bull. 208, 145–155 (2005).

21. Duperron, S. et al. Diversity, relative abundance and metabolic potential of bacterial endosymbionts in three *Bathymodiolus* mussel species from cold seeps in the Gulf of *Mexico*. Environ. Microbiol. 9, 1423–1438 (2007).

22. Wippler, J. et al. Transcriptomic and proteomic insights into innate immunity and adaptations to a symbiotic lifestyle in the gutless marine worm *Olavius algarvensis*. BMC Genomics 17, 942 (2016).

23. Beinart, R. A. et al. Evidence for the role of endosymbionts in regional-scale habitat partitioning by hydrothermal vent symbioses. Proc. Natl. Acad. Sci. USA 109, E3241–E3250 (2012).

24. Jeannin, P. et al. Complexity and complementarity of outer membrane protein A recognition by cellular and humoral innate immunity receptors. Immunity 22, 551–560 (2005).

25. Kawagucci, S. et al. Fluid chemistry in the Solitaire and Dodo hydrothermal fields of the Central Indian Ridge. Geofluids 16, 988–1005 (2016).

26. Ji, F. et al. Geochemistry of hydrothermal vent fluids and its implications for subsurface processes at the active Longqi hydrothermal field, Southwest Indian Ridge. Deep-Sea Res. PT. I 122, 41–47 (2017).

27. Tao, C. et al. Deep high-temperature hydrothermal circulation in a detachment faulting system on the ultra-slow spreading ridge. Nat. Commun. 11, 1300 (2020).

28. Ponnudurai, R. et al. Metabolic and physiological interdependencies in the *Bathymodiolus azoricus* symbiosis. ISME J. 11, 463–477 (2017).

29. Fisher, C. R., Kennicutt, M. C. & Brooks, J. M. Stable carbon isotopic evidence for carbon limitation in hydrothermal vent vestimentiferans. Science 247, 1094–1096 (1990).

30. Rubin-Blum, M. et al. Fueled by methane: deep-sea sponges from asphalt seeps gain their nutrition from methane-oxidising symbionts. ISME J. 13, 1209 (2019).

31. Price, D. R. & Wilson, A. C. A substrate ambiguous enzyme facilitates genome reduction in an intracellular symbiont. BMC Biol. 12, 110 (2014)

32. Kleiner, M., Young, J. C., Shah, M., VerBerkmoes, N. C. & Dubilier, N. Metaproteomics reveals abundant transposase expression in mutualistic endosymbionts. mBio 4, e00223–13 (2013).

33. Wick, R. R., Judd, L. M. & Holt, K. E. Performance of neural network basecalling tools for Oxford Nanopore sequencing. Genome Biol. 20, 129 (2019).

34. Lieberman-Aiden, E. et al. Comprehensive mapping of long-range interactions reveals folding principles of the human genome. Science 326, 289–293 (2009).

35. Bolger, A. M., Lohse, M. & Usadel, B. Trimmomatic: a flexible trimmer for Illumina sequence data. Bioinformatics 30, 2114–2120 (2014).

36. Nurk, S., Meleshko, D., Korobeynikov, A. & Pevzner, P. A. metaSPAdes: a new versatile metagenomic assembler. Genome Res. 27, 824–834 (2017).

37. Wick, R. R., Judd, L. M., Gorrie, C. L. & Holt, K. E. Unicycler: resolving bacterial genome assemblies from short and long sequencing reads. PLoS Comput. Biol. 13, e1005595 (2017).

38. Boetzer, M. & Pirovano, W. SSPACE-LongRead: scaffolding bacterial draft genomes using long read sequence information. BMC Bioinformatics 15, 211 (2014).

39. Parks, D. H., Imelfort, M., Skennerton, C. T., Hugenholtz, P. & Tyson, G. W. CheckM: assessing the quality of microbial genomes recovered from isolates, single cells, and metagenomes. Genome Res. 25, 1043–1055 (2015).

40. Koren, S. et al. Canu: scalable and accurate long-read assembly via adaptive k-mer weighting and repeat separation. Genome Res. 27, 722–736 (2017).

41. Ruan, J. & Li, H. Fast and accurate long-read assembly with wtdbg2. Nat. Methods 17, 155–158 (2019).

42. Vaser, R., Sović, I., Nagarajan, N. & Šikić, M. Fast and accurate *de novo* genome assembly from long uncorrected reads. Genome Res. 27, 737–746 (2017).

43. Walker, B. J. et al. Pilon: an integrated tool for comprehensive microbial variant detection and genome assembly improvement. PloS ONE 9, e112963 (2014).

44. Langmead, B. & Salzberg, S. L. Fast gapped-read alignment with Bowtie 2. Nat. Methods 9, 357–359 (2012).

45. Wu, Y. W., Simmons, B. A. & Singer, S. W. MaxBin 2.0: an automated binning algorithm to recover genomes from multiple metagenomic datasets. Bioinformatics 32, 605–607 (2015).

46. Servant, N. et al. HiC-Pro: An optimized and flexible pipeline for Hi-C processing. Genome Biol. 16, 259 (2015).

47. Durand, N. C. et al. Juicer provides a one-click system for analyzing loop-resolution Hi-C experiments. Cell Syst. 3, 95–98 (2016).

48. Dudchenko, O. et al. *De novo* assembly of the *Aedes aegypti* genome using Hi-C yields chromosome-length scaffolds. Science 356, 92–95 (2017).

49. Simão, F. A., Waterhouse, R. M., Ioannidis, P., Kriventseva, E. V. & Zdobnov, E. M. BUSCO: assessing genome assembly and annotation completeness with single-copy orthologs. Bioinformatics 31, 3210–3212 (2015).

50. Hyatt, D. et al. Prodigal: prokaryotic gene recognition and translation initiation site identification. BMC Bioinformatics 11, 119 (2010).

51. Cantarel, B. L. et al. MAKER: An easy-to-use annotation pipeline designed for emerging model organism genomes. Genome Res. 18, 188–196 (2008).

52. Moriya, Y., Itoh, M., Okuda, S., Yoshizawa, A. C. & Kanehisa, M. KAAS: an automatic genome annotation and pathway reconstruction server. Nucleic Acids Res. 35, W182–W185 (2007).

53. Huerta-Cepas, J. et al. Fast genome-wide functional annotation through orthology assignment by eggNOG-mapper. Mol. Biol. Evol. 34, 2115–2122 (2017).

54. El-Gebali, S. et al. The Pfam protein families database in 2019. Nucleic Acids Res. 47, D427–D432 (2019).

55. Emms, D. M. & Kelly, S. OrthoFinder: solving fundamental biases in whole genome comparisons dramatically improves orthogroup inference accuracy. Genome Biol. 16, 157 (2015).

56. Katoh, K., Kuma, K. I., Toh, H. & Miyata, T. MAFFT version 5: improvement in accuracy of multiple sequence alignment. Nucleic Acids Res. 33, 511–518 (2005).

57. Price, M. N., Dehal, P. S. & Arkin, A. P. FastTree: computing large minimum-evolution trees with profiles instead of a distance matrix. Mol. Biol. Evol. 26, 1641–1650 (2009)

58. dos Reis, M. & Yang, Z. Approximate likelihood calculation on a phylogeny for Bayesian estimation of divergence times. Mol. Biol. Evol. 28, 2161–2172 (2011).

59. Han, M. V., Thomas, G. W., Lugo-Martinez, J. & Hahn, M. W. Estimating gene gain and loss rates in the presence of error in genome assembly and annotation using CAFE 3. Mol. Biol. Evol. 30, 1987–1997 (2013).

60. Nguyen, L. T., Schmidt, H. A., Von Haeseler, A. & Minh, B. Q. IQ-TREE: a fast and effective stochastic algorithm for estimating maximum-likelihood phylogenies. Mol. Biol. Evol. 32, 268–274 (2015).

61. Bray, N. L., Pimentel, H., Melsted, P. & Pachter, L. Near-optimal probabilistic RNA-seq quantification. Nat. Biotechnol. 34, 525 (2016).

62. Robinson, M. D., McCarthy, D. J. & Smyth, G. K. edgeR: a Bioconductor package for differential expression analysis of digital gene expression data. Bioinformatics 26, 139–140 (2010).

63. Wang, Y. et al. MCScanX: a toolkit for detection and evolutionary analysis of gene synteny and collinearity. Nucleic Acids Res. 40, e49–e49 (2012).

64. Wessel, D. M. & Flügge, U. I. A method for the quantitative recovery of protein in dilute solution in the presence of detergents and lipids. Anal Biochem. 138, 141–143 (1984).

