## Supplementary Info for "Dual symbiosis in the deep-sea hydrothermal vent snail *Gigantopelta aegis* revealed by its hologenome"

### Materials and Methods

#### *Genome Survey and Genome Assembly Pipelines*

A 17-mer frequency distribution was obtained using the assemble mode of Platanus version 1.2.4[1] in order to assess the genome characteristics, including genome size, heterozygosity and repetitive content. Both PacBio-only assembly (Canu version 1.7.1[2] correction [genomeSize = 1.27 Gb, corMhapSensitivity = normal, corMinCoverage = 0, corMaxEvidenceErate = 0.15, correctedErrorRate = 0.065, minReadLength = 8000] + SMARTdenovo [-c 1; https://github.com/ruanjue/smartdenovo]; Canu version 1.7.1[2] correction + wtdbg2 version 2.1[3] [-e 2 --tidy-reads 5000 -S 1 -k 15 -p 0 --rescue-low-cov-edges --aln-noskip]; SMARTdenovo [-c 1]; Minimap2 [-x ava-pb] + miniasm[4,5] ; Flye[6]) and hybrid assembler MaSuRCA version 3.2.6[7] pipelines were applied to the genome assembly. In hybrid assembly pipeline, the clean Illumina reads and the Canu corrected subreads were both used. And both the mean and standard deviation of the Illumina library insert size were estimated by Platanus version 1.2.4 assembling[1].

#### *Repeats Annotation*

The species-specific repeats library of the *G. aegis* was *de novo* identified and classified by RepeatModeler version 1.0.11 (http://www.repeatmasker.org/RepeatModeler/) pipeline implemented with RepeatScout version 1.0.5[8], RECON version 1.08[9], TRF version 4.09[10], and NSEG[11]. The repeats of the genome were searched against the species-specific library, RepBase library (RepeatMasker Edition released on 2018, October 26th)[12] as well as Dfam library version 2.0[13] by NCBI RMBlast version 2.6.0, and the hitted regions were further soft-masked by RepeatMasker version 4.0.8 (http://www.repeatmasker.org/RMDownload.html) with a parameter "-xsmall".

#### *Genes Prediction of the Host*

To obtain EST evidence, Trinity version 2.8.3[14] was used to *de novo* assemble and genome-guided assemble the transcripts using the transcriptome sequencing data of all available organs. The *de novo* assembled and genome-guided assembled transcripts were integrated by PASA pipeline version 2.2.0[15] with standalone BLAT version 36x4[16]. The redundancy was removed from the integrated transcripts by using CD-HIT-EST version 4.6.8[17] with "-c 0.95". The non-redundant transcripts served as EST evidence in the gene prediction. To train the model of *ab initio* gene prediction, first round of MAKER was performed with only EST evidence by using "est2genome = 1". The predicted genes of this round of MAKER were only used for training *ab initio* gene prediction model. Only the genes with an annotation edit distance (AED) score equal to zero, with the distances of the neighbouring genes larger than 3 kb, and with more than 3 exons, were used to training Augustus version 3.2.3[18]. After that, MAKER version 2.31.10[19] was performed for a second round to predict genes of the genome with all evidences.

*Mitochondria Genome Assembly and Characteristics*

The mitochondrial genome was assembled by MEGAHIT version 1.1.1[20] using the clean Illumina reads. MITOS Web Sever[21] was used to annotate the protein-coding genes (PCGs), transfer RNA (tRNA) genes, and ribosomal RNA (rRNA) genes of the mitochondrial genome.

*Molecular Clock Analysis*

The following 7 fossil records and geographic events were used to calibrate the phylogenetic tree: minimum = 465.0 Ma for the first appearance of Pteriomorpha[22]; and minimum = 168.6 Ma and soft maximum = 473.4 Ma for *A. californica* and *R. auricularia*[23]; a hard max time-point of 150 Ma for *L. nyassanus and P. canaliculata*, which correspond to the split of South America and Africa[24]; hard minimum bound = 390 Ma for Caenogastropoda and Heterobranchia[25]; minimum = 470.2 Ma and soft maximum = 531.5 Ma for *A. californica* and *L. gigantean*[23], and minimum = 532 Ma and soft maximum = 549 Ma for the first appearance of molluscs[26]; and minimum = 550.25 Ma and soft maximum = 636.1Ma for the first appearance of Lophotrochozoan[26].

*Proteomic Approach*

The oesophageal glands from three *Gigantopelta aegis* were dissected, lysed in a buffer (8M urea, 40mM HEPES, pH=8.0), and sonicated by QSonica (Newtown, CT, USA). The mixtures were centrifuged at 15,000 g for 15min. Methanol-chloroform protein precipitation method[27] was used to purify the supernatant and concentrate the protein yielded. SDS-PAGE (sodium dodecyl sulphate–polyacrylamide gel electrophoresis) gel was used to separate different size of ~30 μg extracted protein, and stained by colloidal coomassie blue. Each gel was cut into 6 slices and dehydrated in an ACN buffer (100mM $NH_4HCO_3$, 50mM $NH_4HCO_3$ and 50% ACN, and 100% ACN). The peptide for LC-MS/MS was obtained through protein reduction (10mM DTT for 45 mim at $56^{o}C$; alkylated by 55mM iodoacetamide for 20 min in dark), protein digestion (sequencing grade Trypsin for 14 hours at $37^{o}C$), peptide extraction (5% formic acid in 50% ACN and 100% ACN sequentially) and further dry (speed-vacuum and desalted with a C18 Sep-Pak column).

Dionex UltiMate 3000 RSLCnano coupled with an Orbitrap Fusion Lumos Mass Spectrometer (Thermo Fisher) was utilised to analyse each protein fraction with the settings: flow rate at 300 nL/min, positive ion mode, 400−1500m/z scan range, 60,000 MS resolution, 1.6m/z isolation window, 40s dynamic exclusion duration, 4.0e5 AGC target, 30% HCD collision energy, and 110 m/z first mass.

The search database contains the protein sequences predicted from the genome and the corresponding reversed sequences (decoy) of both *Gigantopelta aegis* and its two endosymbionts. Mascot version 2.3.0 was used to identify and quantify the protein via the raw mass spectrometry data with settings: 0.6 Da for fragments, 5ppm for precursor, fixed modification: carbamidomethyl (cysteine), variable modification: oxidation (methionine), and up to two missed trypsin cleavage. Peptide with an expectation level smaller than 0.05 was filtered with a false discovery rate of 2.5%.

### Results and Discussion

#### *Genome Survey*

According to 17-mer histogram of *G. aegis* (**Fig. S15**), the genome size was assessed to be 1.21 Gb, the heterozygosity was around 0.5%, and the repeats composition was approximately 50%.

#### *Mitogenome*

A near complete mitochondrial genome of *G. aegis* was assembled into one contig with a length of 16,097 nt (**Fig. S16**). It possesses 37 genes, including 13 protein coding genes, 22 tRNA genes, and 2 rRNA genes (**Table S17**). Twenty genes are in the plus strand, and other 17 ones are in the negative strand. Among the protein coding genes, *nad4l* and *nad4* have a 3 bp overlap.

#### *Assembly, Genes Prediction, and Functional Annotation of Hologenome*

Besides the SOB and the MOB, other bacteria species with low abundance were also found in the sequencing data. It was either contamination from sea water or symbiont that may barely contribute to the symbiosis due to such low abundance. We obtained a total of 4,528,840 Nanopore subreads of 5.77 Gb and 493,104,042 clean Illumina short reads. In the SOB genome, 5,105 genes (92.5%) had hits in the NR database, 3,584 (65.0%) in the GO database, 1,813 (32.9%) in the KEGG database, and 4,405 (79.8%) in the COG database. In the MOB genome, 3,019 (97.3%) genes were annotated in the NR database, 1,969 (63.5%) in the GO database; 1,525 (49.2%) in the KEGG database, and 2,746 (88.5%) were assigned to the COG categories (**Table S4**). The COG annotation and GO annotation of the SOB and the MOB were classified into different functional categories (**Fig. S17, Fig. S18**).

A total of around 121 Gb PacBio raw subreads with an N50 of 10k nt (**Table S14**) and around 171 Gb Illumina raw reads with a length of 150 bp were obtained. Around 52 Gb PacBio corrected subreads with an N50 of 11k nt (**Table S14**) generated 9,479 assembled contigs with an N50 of 471.7 Kb (**Table S16**). A total of 1,607,962,673 pair of Hi-C raw sequencing reads of 482 Gb data generated 56,164,489 pair of valid reads for genome scaffolding (**Table S14**). A total of 5,231 scaffolds with a size of 1.15 Gb and a scaffolds N50 of 81.6 Mb included 15 pseudo-chromosomes (**Table S1**). Among the contigs, 5,216 of them were not anchored into the chromosome groups due to the lack of Hi-C linkage among them or their high repetitiveness (**Table S1**). In the host genome, 50.8 % of the genome were repeats and 74% of these repeats are unclassified (**Fig. S1, Fig. S2, Table S2)**. The Hi-C contact maps of the 15 pseudo-chromosomes were showed in the **Figure S13**. BUSCO (Benchmarking Universal Single-Copy Orthologs) assessment showed the assembled genome had around 94% completeness. Bacteria nucleotides were barely found in the scaffolds of *G. aegis*, suggesting a lack of horizontal gene transfer between the host and the symbionts.

A total of 21,472 genes were predicted from the genome. Among them, 18,507 (86.2%) genes had significant hits in the NCBI NR database, 7,487 (34.9%) in the KEGG database, 13,650 (63.6%) in the KOG database, and 19,165 (89.3%) genes had hits in the functional domains of Pfam database.

*Gene Family Expansion*

Several gene families that are involved in immune recognition were expanded (**Table S11**). Carcinoembryonic antigen-related cell adhesion molecule 5 (CEACAM5) and multiple epidermal growth factor-like domains protein 10 (MEGF) were reported as microbial recognition receptor in oysters[28]. The expansion of these gene families help improve the diversity of microbial recognition patterns, which may help *G. aegis* to recognize its two physiologically distinct types of endosymbionts.

Transposase genes are particularly enriched in bacterial lineages that recently transitioned to a host-associated lifestyle[29,30] and are correspondingly reduced in the ancient, host-restricted bacterial lineages[31]. Transposase enrichment is thus a sign of a recent host-associated lifestyle for the symbionts of *G. aegis*, especially compared to symbionts of *C. squamiferum*. This is corroborated by the fact that the genome size of SOBs in *G. aegis* (4.91 Mb) is much larger than that of the SOBs in *C. squamiferum* (2.59 Mb[32]). Cryptometamorphosis in *Gigantopelta* may be interpreted as an intermediate step before acquiring full reliance on endosymbiosis immediately after settlement, as is the case in *C. squamiferum*. It is possible that *C. squamiferum* also went through a cryptometamorphosis stage sometime during the evolution of the holobiont condition. Since its discovery, evidence has appeared that cryptometamorphosis is more widespread and not unique to *Gigantopelta*. For example, the giant chemosymbiotic shipworm *Kuphus polythalamius* initially settles on wood (like other non-chemosymbiotic shipworms) and only later acquires chemosymbiosis when moving into mud[33]. Such occurrence of cryptometamorphosis in a completely independent chemosymbiotic lineage is suggestive that perhaps a cryptometamorphosis stage is a common 'stepping stone' route towards immediate formation and reliance on symbiosis upon settlement.

*Synteny*

The distribution pattern of non-cross chromosomal synteny was presented for the first time in mollusca and in deep-sea fauna, which may also exist in other closely related groups. These synteny contains genes highly expressed in the oesophageal gland of both *G. aegis* and *C. squamiferum* except the VDG3 gene (**Fig. S19**) that are involved in regulating the development of the digestive system in Mollusca[34,35].

### References


1. Kajitani, R. et al. Efficient *de novo* assembly of highly heterozygous genomes from whole-genome shotgun short reads. *Genome Res.* **24**, 1384–1395 (2014).
2. Koren, S. et al. Canu: scalable and accurate long-read assembly via adaptive k-mer weighting and repeat separation. *Genome Res.* **27**, 722–736 (2017).
3. Ruan, J. & Li, H. Fast and accurate long-read assembly with wtdbg2. *Nat. Methods* **17**,

155–158 (2019).

4. Li, H. Minimap and miniasm: fast mapping and de novo assembly for noisy long sequences. *Bioinformatics* **32**, 2103–2110 (2016).
5. Li, H. Minimap2: pairwise alignment for nucleotide sequences. *Bioinformatics* **34**, 3094–3100 (2018).
6. Kolmogorov, M., Yuan, J., Lin, Y. & Pevzner, P. A. Assembly of long, error-prone reads using repeat graphs. *Nat. Biotechnol.* **37**, 540–546 (2019).
7. Zimin, A. V. et al. The MaSuRCA genome assembler. *Bioinformatics* **29**, 2669–2677. (2013).
8. Price, A. L., Jones, N. C. & Pevzner, P. A. *De novo* identification of repeat families in large genomes. *Bioinformatics* **21**, i351–i358 (2005).
9. Bao, Z. & Eddy, S. R. Automated *de novo* identification of repeat sequence families in sequenced genomes. *Genome Res.* **12**, 1269–1276 (2002).
10. Benson, G. Tandem repeats finder: a program to analyse DNA sequences. *Nucleic Acids Res.* **27**, 573–580 (1999).
11. Wootton, J. C. & Federhen, S. Statistics of local complexity in amino acid sequences and sequence databases. *Comput. Chem.* **17**, 149–163 (1993).
12. Kapitonov, V. V. & Jurka, J. A universal classification of eukaryotic transposable elements implemented in Repbase. *Nat. Rev. Genet.* **9**, 411–412 (2008).
13. Hubley, R. et al. The Dfam database of repetitive DNA families. *Nucleic Acids Res.* **44**, D81–D89 (2015).
14. Haas, B. J. et al. *De novo* transcript sequence reconstruction from RNA-Seq: reference generation and analysis with Trinity. *Nat. Protoc.* **8**, 1494–1512 (2013).
15. Haas, B. J. et al. Improving the *Arabidopsis* genome annotation using maximal transcript alignment assemblies. *Nucleic Acids Res.* **31**, 5654–5666 (2003).
16. Kent, W. J. BLAT—the BLAST-like alignment tool. *Genome Res.* **12**, 656–664 (2002).
17. Li, W. & Godzik, A. Cd-hit: a fast program for clustering and comparing large sets of protein or nucleotide sequences. *Bioinformatics* **22**, 1658–1659 (2006).
18. Stanke, M., Diekhans, M., Baertsch, R. & Haussler, D. Using native and syntenically mapped cDNA alignments to improve *de novo* gene finding. *Bioinformatics* **24**, 637–644 (2008).
19. Cantarel, B. L. et al. MAKER: An easy-to-use annotation pipeline designed for emerging model organism genomes. *Genome Res.* **18**, 188–196 (2008).
20. Li, D., Liu, C. M., Luo, R., Sadakane, K. & Lam, T. W. MEGAHIT: an ultra-fast single-node solution for large and complex metagenomics assembly via succinct *de Bruijn* graph. *Bioinformatics* **31**, 1674–1676 (2015).
21. Bernt, M. et al. MITOS: improved *de novo* metazoan mitochondrial genome annotation. *Mol. Phylogenet. Evol.* **69**, 313–319 (2013).
22. Stöger, I. et al. The continuing debate on deep molluscan phylogeny: evidence for serialia (Mollusca, Monoplacophora + Polyplacophora). *Biomed. Res. Int.* **2013**, 407072 (2013).
23. Benton, M. J., Donoghue, P. C. J. & Asher, R. J. in *The Timetree of Life* (eds. S. Blair Hedges, S. & Kumar, S.) 35–86 (Oxford University Press, 2009).
24. Hayes, K. A. et al. Molluscan models in evolutionary biology: apple snails (Gastropoda: Ampullariidae) as a system for addressing fundamental questions. *Am. Malacol. Bull.* **27**,

47–59 (2009).

25. Jörger, K. M. et al. On the origin of Acochlidia and other enigmatic euthyneuran gastropods, with implications for the systematics of Heterobranchia. *BMC Evol. Biol.* **10**, 323 (2010).
26. Benton, M. J. et al. Constraints on the timescale of animal evolutionary history. *Palaeontol. Electron.* **18**, 1–106 (2015).
27. Wessel, D. M. & Flügge, U. I. A method for the quantitative recovery of protein in dilute solution in the presence of detergents and lipids. *Anal. Biochem.* **138**, 141–143 (1984).
28. Chen, H. et al. The comprehensive immunomodulation of NeurimmiRs in haemocytes of oyster *Crassostrea gigas* after acetylcholine and norepinephrine stimulation. *BMC Genomics* **16**, 942 (2015).
29. Newton, I. L. & Bordenstein, S. R. Correlations between bacterial ecology and mobile DNA. *Curr. Microbiol.* **62**, 198–208 (2011).
30. Kleiner, M., Young, J. C., Shah, M., VerBerkmoes, N. C. & Dubilier, N. Metaproteomics reveals abundant transposase expression in mutualistic endosymbionts. *mBio* **4**, e00223-13 (2013).
31. Moran, N. A., & Plague, G. R. Genomic changes following host restriction in bacteria. *Curr. Opin. Genet. Dev.* **14**, 627–633 (2004).
32. Nakagawa, S. et al. Allying with armored snails: the complete genome of gammaproteobacterial endosymbiont. *ISME J.* **8**, 40–51 (2014).
33. Shipway, J. R. et al. Observations on the life history and geographic range of the giant chemosymbiotic shipworm *Kuphus polythalamius* (Bivalvia: Teredinidae). *Biol. Bull.* **235**, 167–177 (2018).
34. Huang, Z. X. et al. Pyrosequencing of *Haliotis diversicolor* transcriptomes: insights into early developmental molluscan gene expression. *PLoS ONE* **7**, e51279 (2012).
35. He, T. F., Chen, J., Zhang, J., Ke, C. H. & You, W. W. *SARP19* and *vdg3* gene families are functionally related during abalone metamorphosis. *Dev. Genes. Evol.* **224**, 197–207 (2014).
36. Li, Y., Liles, M. R. & Halanych, K. M. Endosymbiont genomes yield clues of tubeworm success. *ISME J.* **12**, 2785–2795 (2018).
37. Li, Y. et al. Genomic adaptations to chemosymbiosis in the deep-sea seep-dwelling tubeworm *Lamellibrachia luymesi*. *BMC Biol.* **17**, 91 (2019).
38. Perez, M. & Juniper, K. Insights into symbiont population structure among three vestimentiferan tubeworm host species at eastern Pacific spreading centres. *Appl. Environ. Microbiol.* **82**, 5197–5205 (2016).
39. Gardebrecht, A. et al. Physiological homogeneity among the endosymbionts of *Riftia pachyptila* and *Tevnia jerichonana* revealed by proteogenomics. *ISME J.* **6**, 766–776 (2012).
40. Goffredi, S. K. et al. Genomic versatility and functional variation between two dominant heterotrophic symbionts of deep-sea *Osedax* worms. *ISME J.* **8**, 908–924 (2014).
41. Yang, Y. et al. Genomic, transcriptomic, and proteomic insights into the symbiosis of deep-sea tubeworm holobionts. *ISME J.* **14**, 135–150 (2019).
42. Ponnudurai, R. et al. Metabolic and physiological interdependencies in the *Bathymodiolus azoricus* symbiosis. *ISME J.* **11**, 463–477 (2017).

43. Sun, J. et al. Adaptation to deep-sea chemosynthetic environments as revealed by mussel genomes. *Nat. Ecol. Evol.* **1**, 121 (2017).
44. Takishita, K. et al. Genomic evidence that methanotrophic endosymbionts likely provide deep-sea *Bathymodiolus* mussels with a sterol intermediate in cholesterol biosynthesis. *Genome Biol. Evol.* **9**, 1148–1160 (2017).
45. Ponnudurai, R. et al. Genome sequence of the sulphur-oxidising *Bathymodiolus thermophilus* gill endosymbiont. *Stand. Genomic Sci.* **12**, 50 (2017).
46. Ikuta, T. et al. Heterogeneous composition of key metabolic gene clusters in a vent mussel symbiont population. *ISME J.* **10**, 990–1001 (2016).
47. Newton, I. L. G. et al. The *Calyptogena magnifica* chemoautotrophic symbiont genome. *Science* **315**, 998–1000 (2007).
48. Kuwahara, H. et al. Reduced genome of the thioautotrophic intracellular symbiont in a deep-sea clam, *Calyptogena okutanii*. *Curr. Biol.* **17**, 881–886 (2007).
49. Sun, J. et al. The scaly-foot snail genome and the ancient origins of biomineralised armour. *Nat. Commun.* **11**, 1657 (2020).
50. Beinart, R. A., Luo, C., Konstantinidis, K., Stewart, F. & Girguis, P. R. The bacterial symbionts of closely related hydrothermal vent snails with distinct geochemical habitats show broad similarity in chemoautotrophic gene content. *Front. Microbiol.* **10**, 1818 (2019).
51. Rubin-Blum, M. et al. Fueled by methane: deep-sea sponges from asphalt seeps gain their nutrition from methane-oxidising symbionts. *ISME J.* **13**, 1209–1225 (2019).
52. Li, Y. et al. Scallop genome reveals molecular adaptations to semi-sessile life and neurotoxins. *Nat. Commun.* **8**, 1721 (2017).
53. Zhang, G. et al. The oyster genome reveals stress adaptation and complexity of shell formation. *Nature* **490**, 49–54 (2012).
54. Belcaid, M. et al. Symbiotic organs shaped by distinct modes of genome evolution in cephalopods. *Proc. Natl. Acad. Sci. USA* **116**, 3030–3035 (2019).
55. Sun, J. et al. Signatures of divergence, invasiveness, and terrestrialization revealed by four apple snail genomes. *Mol. Biol. Evol.* **36**, 1507–1520 (2019).
56. Gerdol, M., Luo, Y.-J., Satoh, N. & Pallavicini, A. Genetic and molecular basis of the immune system in the brachiopod *Lingula anatina*. *Dev. Comp. Immunol.* **82**, 7–30 (2018).
57. Simakov, O. et al. Insights into bilaterian evolution from three spiralian genomes. *Nature* **493**, 526–531 (2012).
58. Albertin, C. B. et al. The octopus genome and the evolution of cephalopod neural and morphological novelties. *Nature* **524**, 220–224 (2015).
59. Luo, Y. J. et al. Nemertean and phoronid genomes reveal lophotrochozoan evolution and the origin of bilaterian heads. *Nat. Ecol. Evol.* **2**, 141–151 (2018).
60. Takeuchi, T. et al. Bivalve-specific gene expansion in the pearl oyster genome: implications of adaptation to a sessile lifestyle. *Zool. Lett.* **2**, 3 (2016).
61. Wang, S. et al. Scallop genome provides insights into evolution of bilaterian karyotype and development. *Nat. Ecol. Evol.* **1**, 120 (2017).
62. Schell, T. et al. An annotated draft genome for *Radix auricularia* (Gastropoda, Mollusca). *Genome Biol. Evol.* **9**, 585–592 (2017).
63. Masonbrink, R. E. et al. An annotated genome for *Haliotis rufescens* (red abalone) and

resequenced green, pink, pinto, black, and white abalone species. *Genome Biol. Evol.* **11**, 431–438 (2019).

### Tables

**Table S1** Summary of the assembly statistics and functional annotation of *Gigantopelta aegi*s genome. (NR: non-redundant RefSeq protein database; GO: gene ontology database; KEGG: Kyoto encyclopedia of genes and genomes database; KOG: EuKaryotic Orthologous Groups)

| Assembly Feature | Statistics |
|---|---|
| Estimated genome size (17-mer analysis) | 1.21 Gb |
| Number of scaffolds | 5,231 |
| Total assembly size (bp) | 1,149,607,922 |
| Longest scaffolds (nt) | 120,622,278 |
| N50 scaffold length (nt) | 81,591,406 |
| L50 scaffold count | 6 |
| N50 contig length (nt) | 461,769 |
| L50 contig count | 671 |
| Number of contigs | 9,479 |
| **Functional Annotation** | |
| Gene number predicted | 21,472 |
| NR | 19,312 (89.9%) |
| GO | 5,901 (27.5%) |
| KEGG | 7,487 (34.9%) |
| KOG | 13,560 (63.2%) |
| Pfam | 19,165 (89.3%) |
| 15 pseudo-chromosomes (1.006 Gb) + 5,216 contigs (143.6 Mb) | |

**Table S2** Classification and composition of repeats content in the genome of *Gigantopelta aegis* gastropod.

| Repeats | Count | Length occupied | Percentage |
|---|---|---|---|
| DNA transposons | 822 | 242470 | 0.02% |
| Academ | 2581 | 914211 | 0.08% |
| CMC-Chapaev-3 | 222 | 153341 | 0.01% |
| Crypton | 592 | 181221 | 0.02% |
| Ginger | 751 | 227020 | 0.02% |
| IS | 1 | 61 | 0.00% |
| Kolobok-Hydra | 752 | 875613 | 0.08% |
| Maverick | 5332 | 3187177 | 0.28% |
| MuLE-MuDR | 297 | 142706 | 0.01% |
| P | 1711 | 1179122 | 0.10% |
| PIF-Harbinger | 823 | 312158 | 0.03% |
| PIF-ISL2EU | 1091 | 461501 | 0.04% |
| Sola | 4361 | 384141 | 0.03% |
| TcMar-Mariner | 426 | 88052 | 0.01% |
| TcMar-Pogo | 1887 | 753494 | 0.07% |
| TcMar-Tc1 | 26585 | 21697335 | 1.89% |
| hAT-Ac | 2428 | 360679 | 0.03% |
| hAT-Tip100 | 1604 | 306676 | 0.03% |
| LINE | 1921 | 793302 | 0.07% |
| CR1-Zenon | 6990 | 1776404 | 0.15% |
| I | 38938 | 5604071 | 0.49% |
| I-Nimb | 29595 | 9949494 | 0.87% |
| Jockey | 1618 | 1149642 | 0.10% |
| L1 | 2516 | 543469 | 0.05% |
| L1-Tx1 | 11441 | 6333774 | 0.55% |
| L2 | 1477 | 526978 | 0.05% |
| Penelope | 21146 | 5362853 | 0.47% |
| Proto2 | 714 | 407527 | 0.04% |
| RTE-BovB | 2185 | 457991 | 0.04% |
| RTE-X | 22943 | 8511840 | 0.74% |
| **LTR** | | | |
| Copia | 250 | 127009 | 0.01% |
| DIRS | 4136 | 3573205 | 0.31% |
| Gypsy | 14622 | 13102080 | 1.14% |
| Gypsy-Troyka | 156 | 259429 | 0.02% |
| Ngaro | 249 | 157130 | 0.01% |
| Low_complexity | 49932 | 3156496 | 0.28% |
| Simple_repeat | 653717 | 47311156 | 4.12% |
| Unknown | 2333786 | 442290191 | 38.55% |

| | | | |
|---|---|---|---|
| **Total** | **3250598** | **582861019** | **50.80%** |

**Table S4** Genome assembly results and functional annotation results of the sulphur-oxidising bacteria (SOB) and the methane-oxidising bacteria (MOB) housed in the oesophageal gland of *Gigantopelta aegis* (NR: non-redundant RefSeq protein database; GO: gene ontology database; KEGG: Kyoto encyclopedia of genes and genomes database; COG: clusters of orthologous groups).

| | SOB | MOB |
|---|---|---|
| **Genome Assembly** | | |
| Genome Size | 4.91 Mb | 2.93 Mb |
| Contigs Number | 18 | 28 |
| Scaffolds Number | 11 | 10 |
| Contamination | 3.25% | 1.67% |
| Completeness | 98.55% | 99.25% |
| Coverage | ~ 3,050 | ~410 |
| **Functional Annotation** | | |
| Gene Number | 5,518 | 3,102 |
| NR | 5,105 (92.5%) | 3,019 (97.3%) |
| GO | 3,584 (65.0%) | 1,969 (63.5%) |
| KEGG | 1,813 (32.9%) | 1,525 (49.2%) |
| COG | 4,405 (79.8%) | 2,746 (88.5%) |

**Table S10** Available genomes of symbionts belonging to Gammaproteobacteria in deep-sea invertebrate taxa.

| Species | Host | SOB | MOB | Sampling Habitat | NCBI Accession | Reference |
|---|---|---|---|---|---|---|
| Annelida | | | | | | |
| *Escarpia spicata* | | + | | deep-sea seep | QFXE00000000 | Ref 36 |
| *Lamellibrachia luymesi* | + | + | | deep-sea seep | SDWI00000000, QFXD00000000 | Ref 36, 37 |
| *Galathealinum brachiosum* | | + | | deep-sea muddy sediments | QFXC00000000 | Ref 36 |
| *Ridgeia piscesae* | | + | | deep-sea vent | LDXT00000000 | Ref 38 |
| *Riftia pachyptila* | | + | | deep-sea vent | AFOC00000000 | Ref 39 |
| *Seepiophila jonesi* | | + | | deep-sea seep | QFXF00000000 | Ref 36 |
| *Tevnia jerichonana* | | + | | deep-sea vent | AFZB00000000 | Ref 39 |
| *Osedax frankpressi* | | + | | deep-sea whale fall | ASZJ00000000 | Ref 40 |
| *Paraescarpia echinospica* | | + | | deep-sea seep | RZUD00000000 | Ref 41 |
| Mollusca | | | | | | |
| *Bathymodiolus azoricus* | | + | + | deep-sea vent | CDSC00000000, FMJP00000000 | Ref 42 |
| *Bathymodiolus platifrons* | + | | + | deep-sea seep | MJUT00000000.1, PRJDB5337 | Ref 43, 44 |
| *Bathymodiolus thermophilus* | | + | | deep-sea vent | CP024634.1 | Ref 45 |
| *Bathymodiolus* sp. | | | + | deep-sea vent | FNVW00000000 | Ref 42 |
| *Bathymodiolus puteoserpentis* | | + | + | deep-sea vent | FQTQ00000000, UEXF00000000 | - |
| *Bathymodiolus brooksi* | | + | | deep-sea vent | FQTS00000000 | - |
| *Bathymodiolus heckerae* | | + | | deep-sea vent | FXLV00000000 | - |
| *Bathymodiolus septemdierum* | | + | | deep-sea vent | AP013042.1 | Ref 46 |
| *Calyptogena magnifica* | | + | | deep-sea vent | JARW00000000 | Ref 47 |
| *Phreagena okutanii* | | + | | deep-sea vent | AP009247.1 | Ref 48 |
| *Chrysomallon squamiferum* | + | + | | deep-sea vent | PRJNA523462, AP012978.1 | Ref 32, 49 |
| *Alviniconcha* | | + | | deep-sea vent | RAST: 6666666.293770, 6666666.293769 | Ref 50 |
| *Ifremeria nautilei* | | + | + | deep-sea vent | RAST: 6666666.293767, 6666666.296237 | Ref 50 |
| *Gigantopelta aegis* | + | + | + | deep-sea vent | PRJNA612619 | present study |
| Porifera | | | | | | |
| *Iophon methanophila* | | | + | deep-sea seep | PRJNA475442 | Ref 51 |
| *Hymedesmia (Stylopus) methanophila* | | | + | deep-sea seep | PRJNA475438 | |

**Table S11** The function of gene family that are expanded in the *Gigantopelta aegis* genome.

| Description | Gene Family | Function | Corrected P Value |
|---|---|---|---|
| 5-hydroxytryptamine receptor 4 | HTR4 | Immune | 7.32E-04 |
| BTB/POZ domain-containing protein 6 | BTBD6 | Suppress the programmed cell death in innate immune response (Orosa et al. 2017) | 8.10E-10 |
| Carcinoembryonic antigen-related cell adhesion molecule 5 | CEACAM5 | Immune intracellular receptors (Chen et al. 2015) | 7.46E-12 |
| E3 ubiquitin-protein ligase TRIM56 | TRIM | Immune signaling molecules (Chen et al. 2015) | 3.17E-04 |
| E3 ubiquitin-protein ligase rnf213-alpha | E3 | Immune signaling molecules (Chen et al. 2015) | 1.17E-04 |
| Inhibitor of apoptosis-like protein | IAP | Regulation of innate immunity, inflammation and apoptosis (Berthelet & Dubrez 2013) | 6.64E-59 |
| KRAB-A domain-containing protein 2 | KRBA2 | Epigenetic control of adaptive immune (Santoni de Sio et al. 2014) | 9.97E-16 |
| F-Lectin | F-Lectin | Recognition (Innate immunity against microbial invasion; Chen et al. 2011) | 5.01E-07 |
| Multiple epidermal growth factor-like domains protein 10 | MEGF | Recognition (Perovic-Ottstadt et al 2004) | 3.33E-03 |
| Ankyrin repeat protein | ANK | Symbiosis factors (Thomas et al. 2010) | 2.59E-02 |

**Table S12** KEGG enrichment of highly expressed genes in the oesophageal glands of *Gigantopelta aegis*.

| KEGG BRITE ID | Biological Modules | OsG HEG Number | Total Number | Corrected *P* Value |
|---|---|---|---|---|
| ko02000 | Transporters | 36 | 256 | 5.75E-06 |
| ko04147 | Exosome | 35 | 311 | 5.50E-04 |
| ko01000 | Enzymes | 152 | 2071 | 4.38E-04 |
| ko04121 | Ubiquitin system | 10 | 416 | 4.87E-03 |
| ko01002 | Peptidases | 23 | 221 | 7.65E-03 |
| ko03036 | Chromosome and associated proteins | 23 | 663 | 1.30E-02 |
| ko04090 | Cell differenciation molecules | 8 | 55 | 2.66E-02 |
| ko04091 | Lectins | 5 | 32 | 5.81E-02 |
| ko01001 | Protein kinases | 7 | 232 | 6.59E-02 |
| ko01003 | Glycosyltransferases | 8 | 82 | 8.85E-02 |
| **KEGG Pathway ID** | **Pathway** | | | |
| 04142 | Lysosome | 22 | 76 | 7.52E-06 |
| 00511 | Other glycan degradation | 6 | 11 | 7.83E-03 |
| 04973 | Carbohydrate digestion and absorption | 5 | 12 | 7.35E-02 |

**Table S14** Pacific Biosciences sequencing data and Hi-C sequencing data of *Gigantopelta aegis*.

| | Size (Gb) | N50 (nt) | Longest (nt) | Reads Number |
|---|---|---|---|---|
| PacBio Subreads | 121 | 10 K | 109 K | 17,718,439 |
| PacBio Corrected Subreads | 52 | 11 K | 109 K | 4,701,940 |
| Hi-C raw reads | 482 | - | - | 3,215,925,346 |
| Hi-C valid reads | - | - | - | 112,328,978 |

**Table S15** Transcriptome sequencing data of four individuals (Ga01 [male], Ga02 [female], Ga03 [female], Ga04 [female]) with different dissected tissues. (meta: metatranscriptome sequencing). The RNA of oesophageal gland was used to construct both eukaryotic library and bacterial library.

| Tissues in different individuals | Raw reads | Clean Reads |
|---|---|---|
| Ga01 Cephalic tentacles | 39,289,210 | 37,854,692 |
| Ga01 Ctenidium | 40,755,872 | 39,275,400 |
| Ga01 Digestive gland | 39,928,094 | 38,574,096 |
| Ga01 Foot | 38,047,576 | 36,749,336 |
| Ga01 Gonad | 42,979,606 | 41,510,252 |
| Ga01 Mantle internal | 40,992,658 | 39,499,540 |
| Ga01 Mantle edge | 38,131,842 | 36,773,324 |
| Ga01 Oesophageal gland | 136,226,224 | 133,303,450 |
| Ga01 Oesophageal gland (meta) | 67,484,722 | 65,380,774 |
| Ga02 Cephalic tentacles | 47,429,082 | 46,646,900 |
| Ga02 Ctenidium | 114,776,966 | 113,018,798 |
| Ga02 Digestive gland | 46,056,578 | 45,353,184 |
| Ga02 Epipodial tentacles | 46,924,072 | 46,101,318 |
| Ga02 Foot | 51,564,116 | 50,777,676 |
| Ga02 Mantle | 62,871,456 | 61,599,812 |
| Ga02 Nephridium | 43,485,296 | 42,859,728 |
| Ga02 Oesophageal gland | 101,244,992 | 99,021,890 |
| Ga02 Operculum | 43,453,604 | 42,782,788 |
| Ga02 Testis | 44,804,546 | 44,040,958 |
| Ga03 Auricle heart | 40,209,102 | 38,544,626 |
| Ga03 Ventricle heart | 61,665,090 | 58,928,864 |
| Ga03 Cephalic tentacles | 38,765,348 | 37,040,658 |
| Ga03 Ctenidium | 42,849,664 | 40,826,230 |
| Ga03 Epipodial tentacles | 40,563,874 | 38,752,620 |
| Ga03 Foot | 38,479,932 | 36,842,512 |
| Ga03 Ovary | 45,663,166 | 44,454,748 |
| Ga03 Mantle internal | 35,283,212 | 33,810,720 |
| Ga03 Mantle edge | 40,839,008 | 39,100,904 |
| Ga03 Oesophageal gland | 263,318,982 | 257,059,132 |
| Ga03 Oesophageal gland (meta) | 71,018,768 | 67,281,746 |
| Ga04 Cephalic tentacles | 45,453,406 | 42,340,348 |
| Ga04 Ctenidium | 41,484,232 | 39,704,082 |
| Ga04 Digestive gland | 50,399,428 | 48,287,006 |
| Ga04 Epipodial tentacles | 41,337,662 | 38,922,138 |
| Ga04 Foot | 43,180,126 | 40,837,562 |
| Ga04 Ovary | 40,721,278 | 38,950,644 |
| Ga04 Mantle edge | 40,583,218 | 38,694,406 |

| | | |
|---|---|---|
| Ga04 Oesophageal gland | 158,304,352 | 153,983,134 |
| Ga04 Oesophageal gland (meta) | 77,014,174 | 73,210,926 |
| Ga04 Ventricle heart | 39,465,714 | 37,728,210 |

**Table S16** Comparison of genome assembly by different assembling pipelines.

| Assembler | Total size | Contig number | N50 | Longest contigs |
|---|---|---|---|---|
| Canu correction+wtdbg2 | 1.16 Gb | 9,842 | 467 kb | 5.4 Mb |
| Canu correction+ SMARTdenovo | 1.12 Gb | 7,995 | 260 Kb | 1.8 Mb |
| SMARTdenovo | 1.31 Gb | 8,520 | 313 Kb | 2.59 Mb |
| Minimap2+miniasm | 1.48 Gb | 18,745 | 183 Kb | 1.38 Mb |
| Flye | 1.26 Gb | 13,214 | 292 Kb | 2.63 Mb |
| Hybrid assembly (MaSuRca) | 1.38 Gb | 18,264 | 202 Kb | 2.76 Mb |

**Table S17** Characterization of *Gigantopelta aegis* mito-genome.

| Gene name | Feature | Start | End | Length | Intergenic nucleotides (nt) | Start codon | Stop codon | Strand |
|---|---|---|---|---|---|---|---|---|
| *trnM(atg)* | tRNA | 444 | 510 | 66 | 443 | | | + |
| *rrnS* | rRNA | 509 | 1386 | 877 | -1 | | | + |
| *trnV(gta)* | tRNA | 1379 | 1443 | 64 | -7 | | | + |
| *rrnL* | rRNA | 1424 | 2803 | 1379 | -19 | | | + |
| *trnL1(cta)* | tRNA | 2779 | 2843 | 64 | -24 | | | + |
| *trnL2(tta)* | tRNA | 2854 | 2920 | 66 | 11 | | | + |
| *nad1* | gene | 2921 | 3826 | 905 | 1 | ATG | CTA | + |
| *trnP(cca)* | tRNA | 3866 | 3932 | 66 | 40 | | | + |
| *nad6* | gene | 3934 | 4419 | 485 | 2 | ATG | CGG | + |
| *trnE(gaa)* | tRNA | 4435 | 4500 | 65 | 16 | | | + |
| *cob* | gene | 4521 | 5639 | 1118 | 21 | ATA | ATT | + |
| *trnS2(tca)* | tRNA | 5652 | 5716 | 64 | 13 | | | + |
| *nad4l* | gene | 5725 | 6021 | 296 | 9 | ATA | TGC | + |
| *nad4* | gene | 6018 | 7385 | 1367 | -3 | ATG | TGA | + |
| *trnH(cac)* | tRNA | 7400 | 7467 | 67 | 15 | | | + |
| *nad5* | gene | 7510 | 9183 | 1673 | 43 | ATC | GTC | + |
| *cox2* | gene | 9207 | 9872 | 665 | 24 | ATG | TGA | + |
| *trnD(gac)* | tRNA | 9898 | 9963 | 65 | 26 | | | + |
| *atp8* | gene | 9964 | 10125 | 161 | 1 | ATG | TGA | + |
| *atp6* | gene | 10213 | 10812 | 599 | 88 | ATT | CAT | + |
| *trnT(aca)* | tRNA | 10861 | 10930 | 69 | 49 | | | - |
| *trnF(ttc)* | tRNA | 10952 | 11019 | 67 | 22 | | | - |
| *cox1* | gene | 11082 | 12593 | 1511 | 63 | ATG | GAG | - |
| *nad2* | gene | 12715 | 13623 | 908 | 122 | ATA | CCT | - |
| *trnS1(agc)* | tRNA | 13679 | 13745 | 66 | 56 | | | - |
| *nad3* | gene | 13759 | 14100 | 341 | 14 | ATT | TGA | - |
| *trnI(atc)* | tRNA | 14104 | 14172 | 68 | 4 | | | - |
| *trnN(aac)* | tRNA | 14173 | 14238 | 65 | 1 | | | - |
| *trnR(cgt)* | tRNA | 14241 | 14310 | 69 | 3 | | | - |
| *trnA(gca)* | tRNA | 14319 | 14385 | 66 | 9 | | | - |
| *trnK(aaa)* | tRNA | 14396 | 14464 | 68 | 11 | | | - |
| *cox3* | gene | 14479 | 15231 | 752 | 15 | GTG | TCT | - |
| *trnG(gga)* | tRNA | 15260 | 15325 | 65 | 29 | | | - |
| *trnY(tac)* | tRNA | 15338 | 15404 | 66 | 13 | | | - |
| *trnC(tgc)* | tRNA | 15405 | 15466 | 61 | 1 | | | - |
| *trnW(tga)* | tRNA | 15467 | 15533 | 66 | 1 | | | - |
| *trnQ(caa)* | tRNA | 15536 | 15608 | 72 | 3 | | | - |

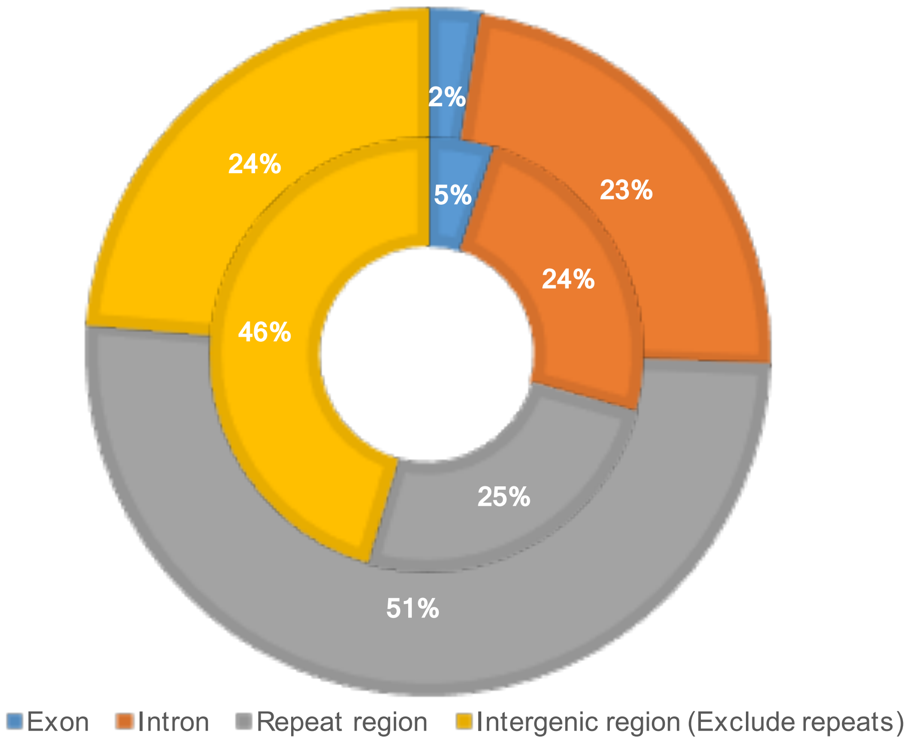


**Figure S1** Composition of different genomic components of *Gigantopelta aegis* and *Chrysomallon squamiferum*. Outer ring: *G. aegis*; Inner ring: *C. squamiferum*.

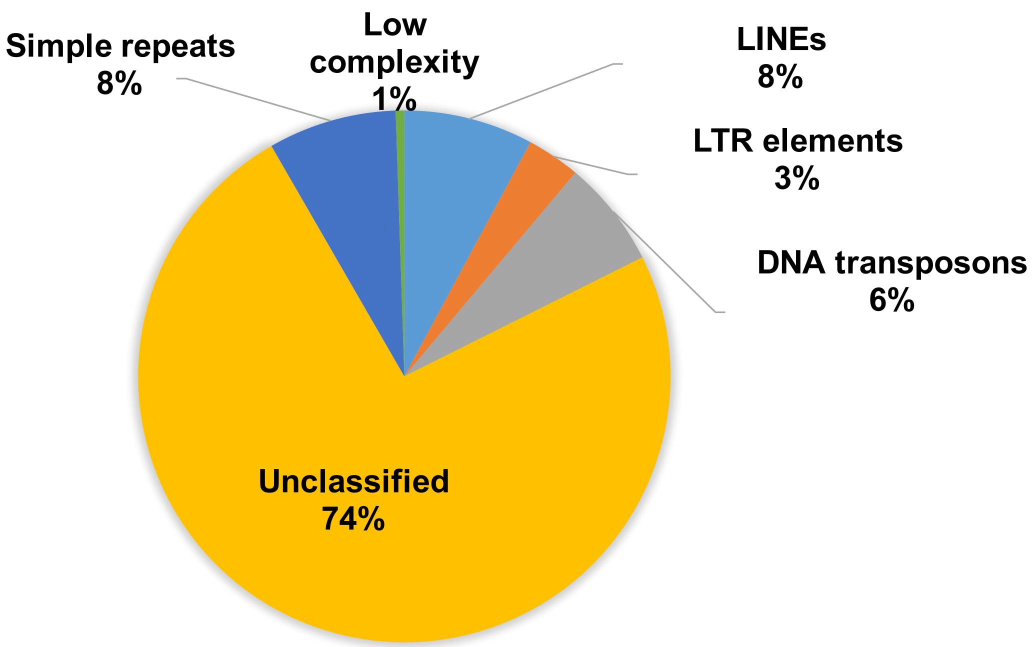


**Figure S2** The repeat component and composition of *Gigantopelta aegis* genome.

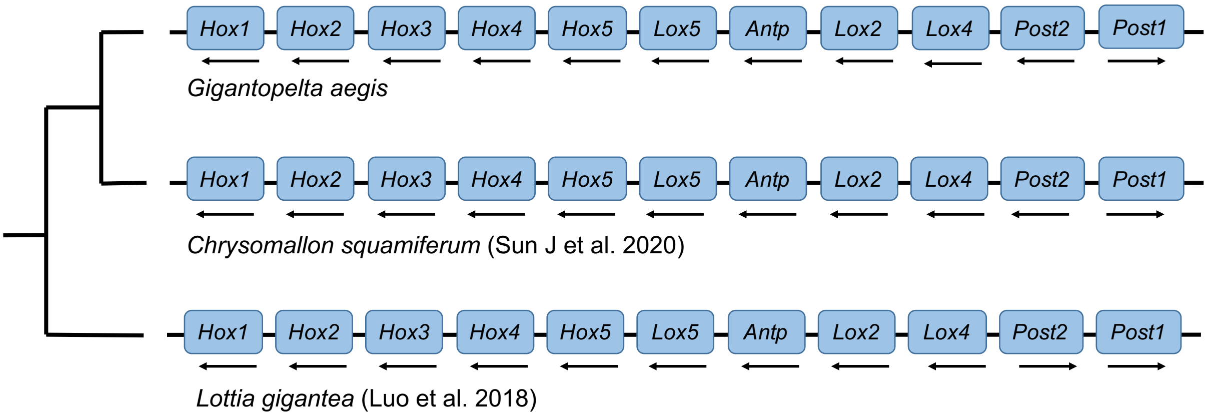


**Figure S3** The hox clusters shared same gene order between *Gigantopelta aegis* and *Chrysomallon squamiferum*.

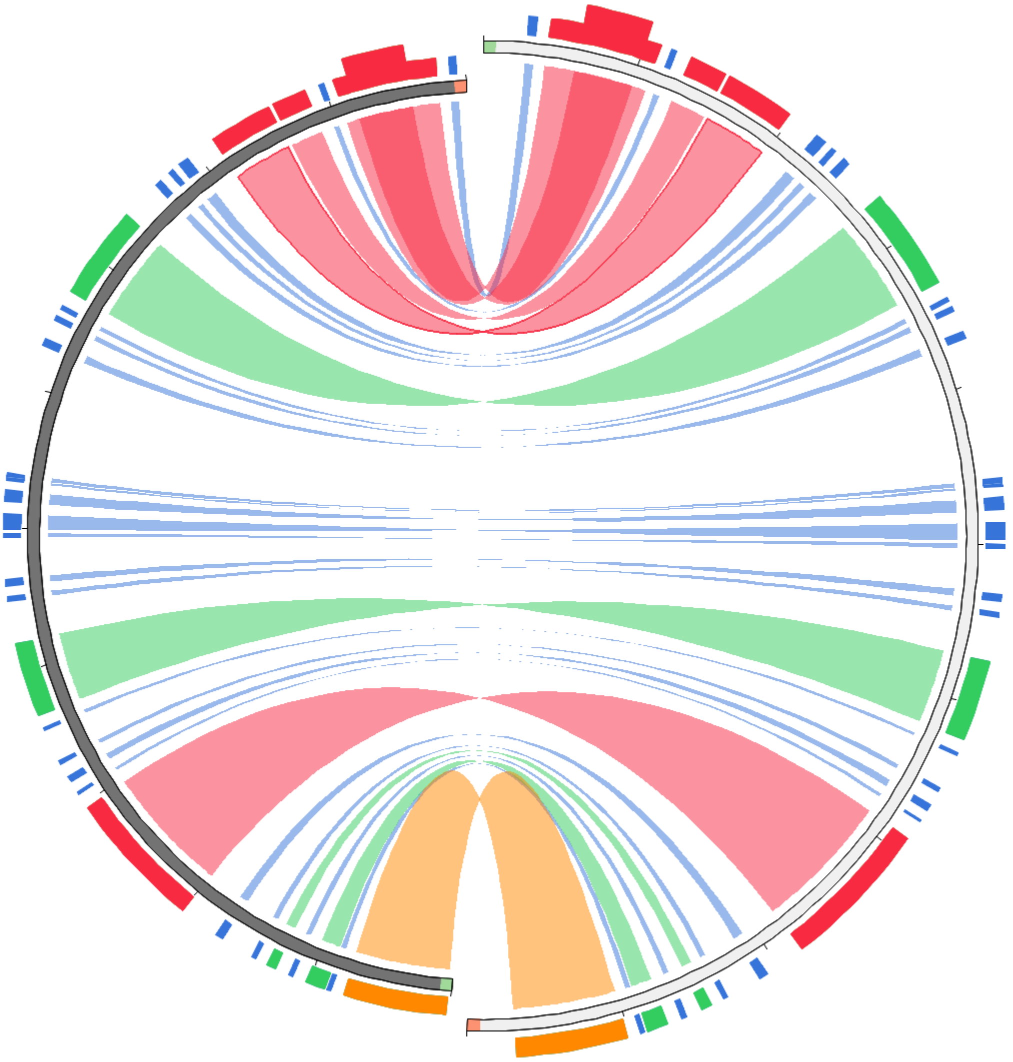

**Figure S4** The synteny of mitochondrial genomes shared between *Gigantopelta aegis* (right) and *Chrysomallon squamiferum* (left).

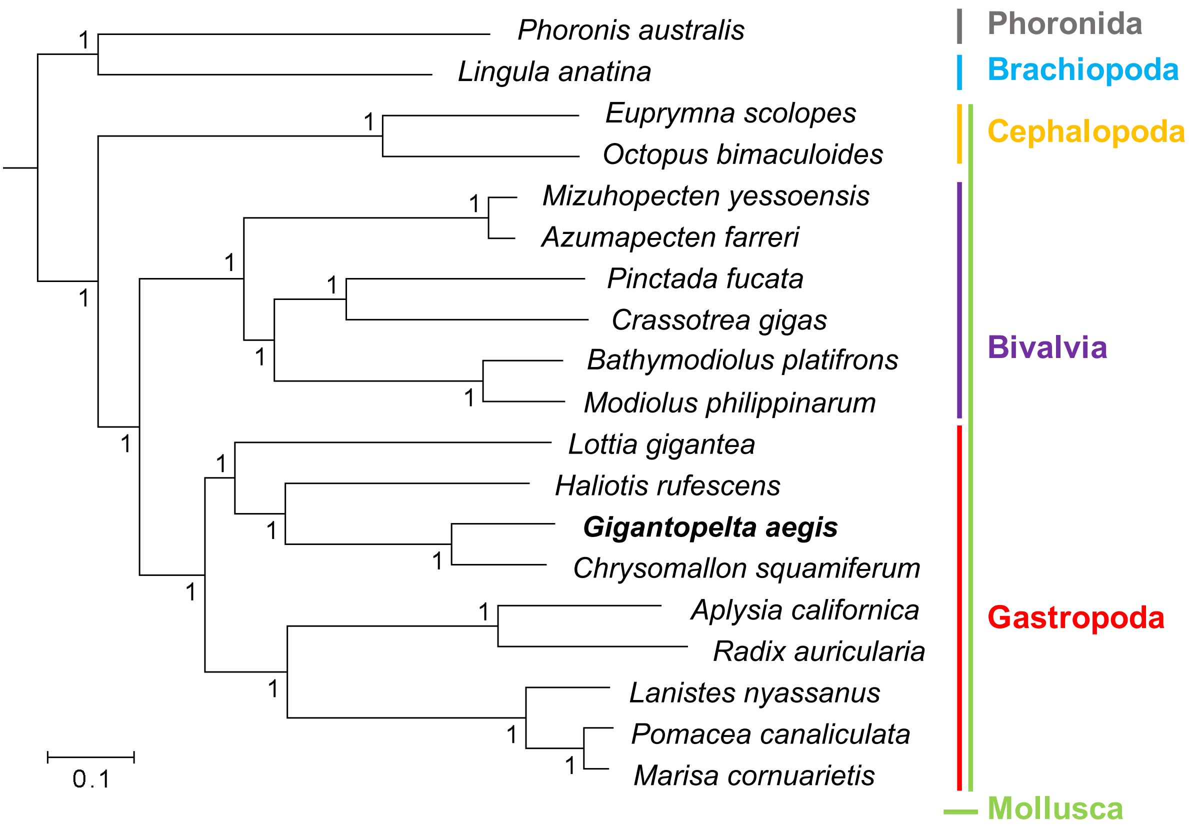


**Figure S5** A phylogenetic tree of *Gigantopelta aegis* and other lophotrochozoan references. The references are listed as follows: *Aplysia californica* (GenBank No. GCA_000002075), *Chrysomallon squamiferum*[49], *Bathymodiolus platifrons*[43], *Modiolus philippinarum*[43], *Azumapecten farreri*[52], *Crassotrea gigas*[53], *Euprymna scolopes*[54], *Lanistes nyassanus*[55], *Marisa cornuarietis*[55], *Pomacea canaliculata*[55], *Lingula anatine*[56], *Lottia gigantea*[57], *Octopus bimaculoides*[58], *Phoronis australis*[59], *Pinctada fucata*[60], *Mizuhopecten yessoensis*[61], *Radix auricularia*[62], *and Haliotis rufescens*[63]).

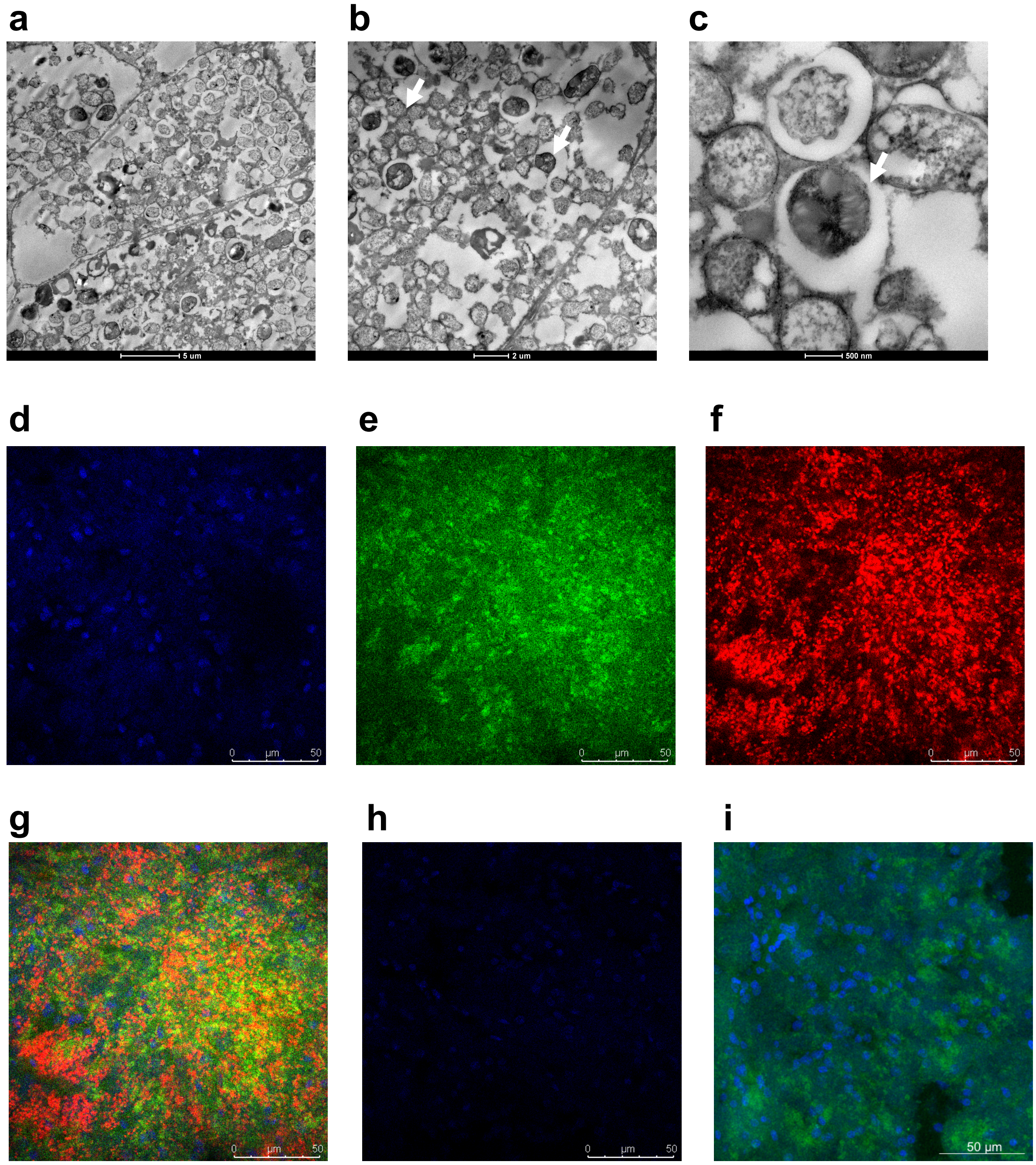


**Figure S6** Transmission electron microscopy (TEM) and fluorescence *in situ* hybridisation (FISH) images of the oesophageal gland from *Gigantopelta aegis*. The TEM images of **a**. an entire bacteriocyte cell housing intracellular endosymbiont (scale bar: 5 µm), **b**. endosymbionts showing two distinct morphological types (white arrows; scale bar: 2 µm), and **c**. one endosymbiont showing intracellular stacked membranes (white arrows; scale bar: 500 nm). FISH images of **d**. host nuclear DNA, **e**. sulphur-oxidising bacteria (SOB), **f**. methane-oxidising bacteria (MOB), and **g**. the merged signal of **d**., **e**., and **f**. on transverse sections of oesophageal gland from *Gigantopelta aegis* (scale bar: 50 µm). FISH experiments were performed with specific 16S rRNA probes for SOB and MOB. FISH image of **h**. Negative control: DNA (DAPI staining) and NON338 probe (Wallner et al. 1993). The signal of

NON338 is bare. **i**. Positive control: DNA (DAPI staining) and universal EUB338 probe (Amann et al. 1990): green signal. Colour: DNA (DAPI staining): blue; SOB (Cy3): green; MOB (Cy5): red.

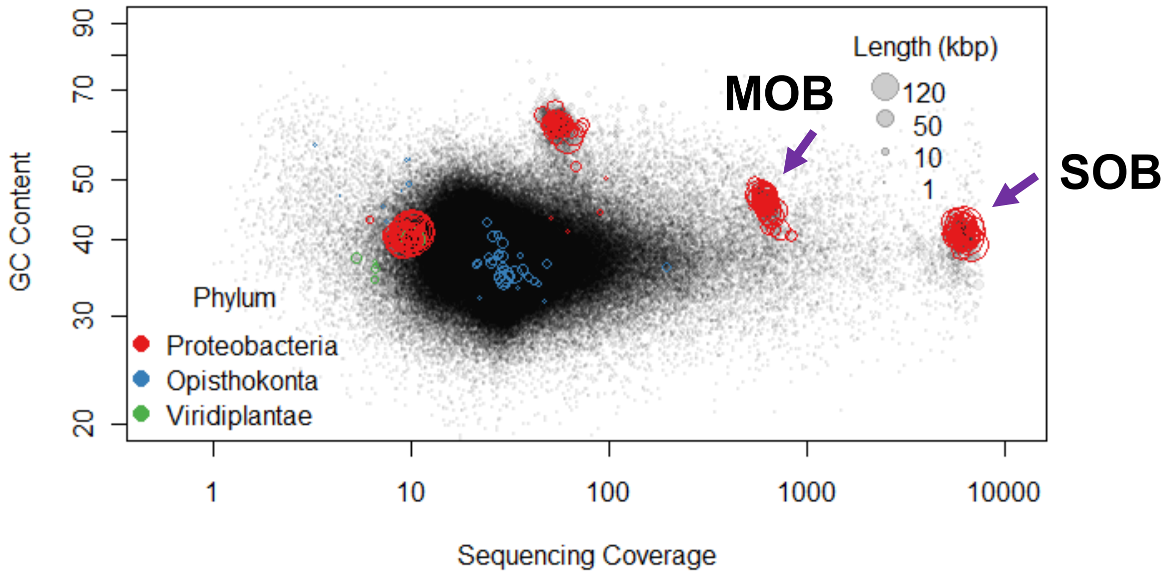


**Figure S7** Genome binning of the initially assembled contigs from the oesophageal gland of *Gigantopelta aegis*. Two dominant bacteria had significantly higher coverage than the host. Each dot represents a contig.

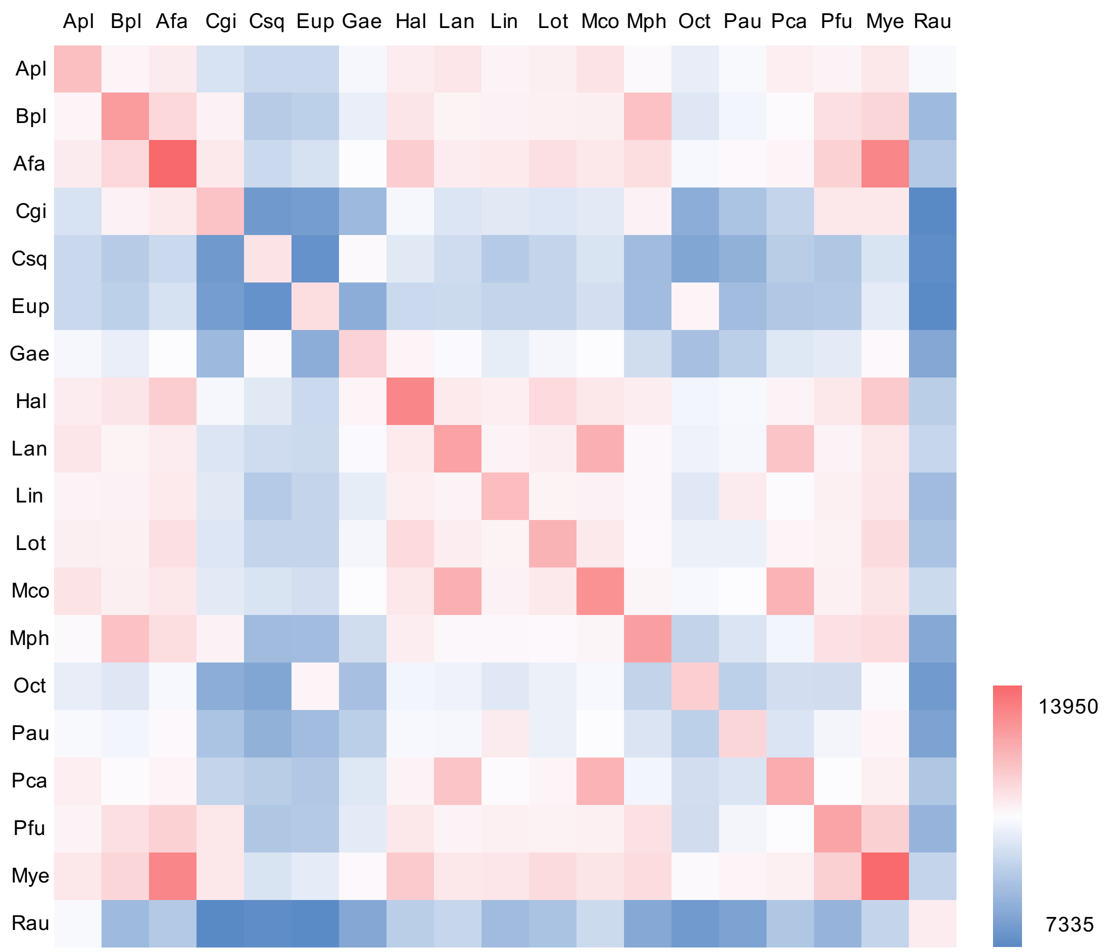


**Figure S8** Heat map of shared gene family numbers of *Gigantopelta aegis* (Gae) and other references (Apl: *Aplysia californica* [GenBank No. GCA_000002075]; Bpl: *Bathymodiolus platifrons*[43]; Afa: *Azumapecten farreri*[52]; Cgi: *Crassotrea gigas*[53]; Csq: *Chrysomallon squamiferum*[49]; Eup: *Euprymna scolopes*[54]; Hal: *Haliotis rufescens*[63]; Lan: *Lanistes nyassanus*[55]; Lin: *Lingula anatine*[56]; Lot: *Lottia gigantea*[57]; Mco: *Marisa cornuarietis*[55]; Mph: *Modiolus philippinarum*[43]; Oct: *Octopus bimaculoides*[58]; Pau: *Phoronis australis*[59]; Pca: *Pomacea canaliculata*[55]; Pfu: *Pinctada fucata*[60]; Mye: *Mizuhopecten yessoensis*[61]; Rau: *Radix auricularia*[62]).

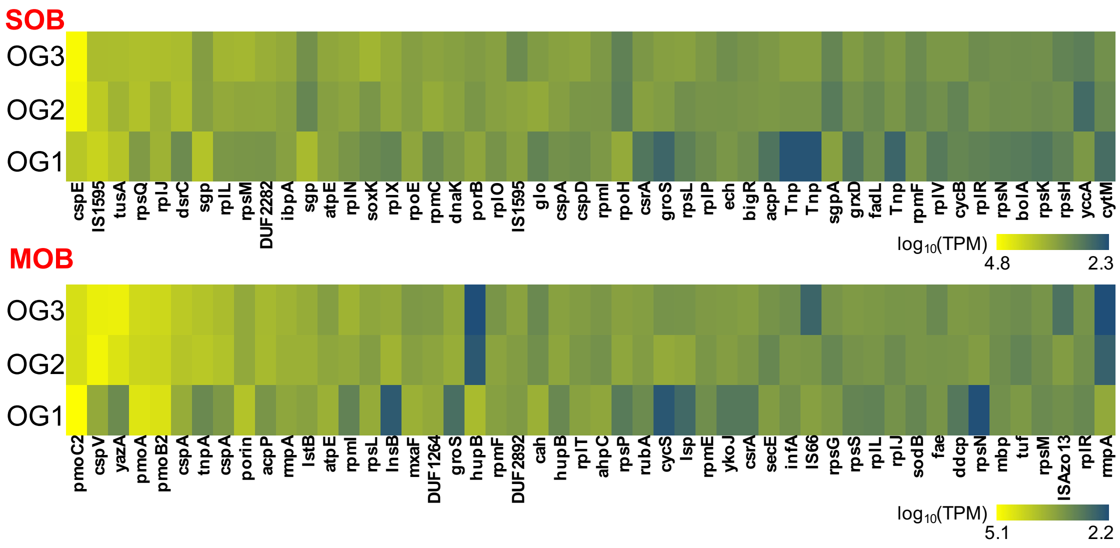


**Figure S9** Genes with top 50 highest gene expression in the sulphur-oxidising symbionts (SOB) and the methane-oxidising symbionts (MOB) of *Gigantopelta aegis* (oesophageal glands from three individuals: OG1, OG2, and OG3; TPM: transcripts per million).

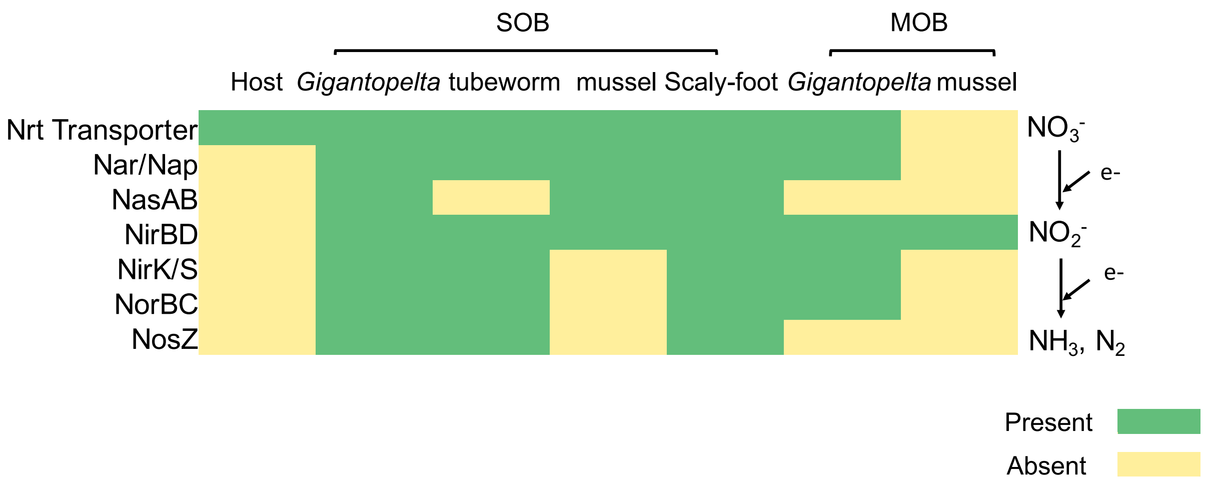


**Figure S10** The nitrate metabolism in the sulphur-oxidising endosymbiont (SOB), the methane-oxidising endosymbiont (MOB), and *Gigantopelta aegis* host, in comparison with the sulphur-oxidising endosymbionts and the methane-oxidising endosymbionts housed in tube worms and mussels.

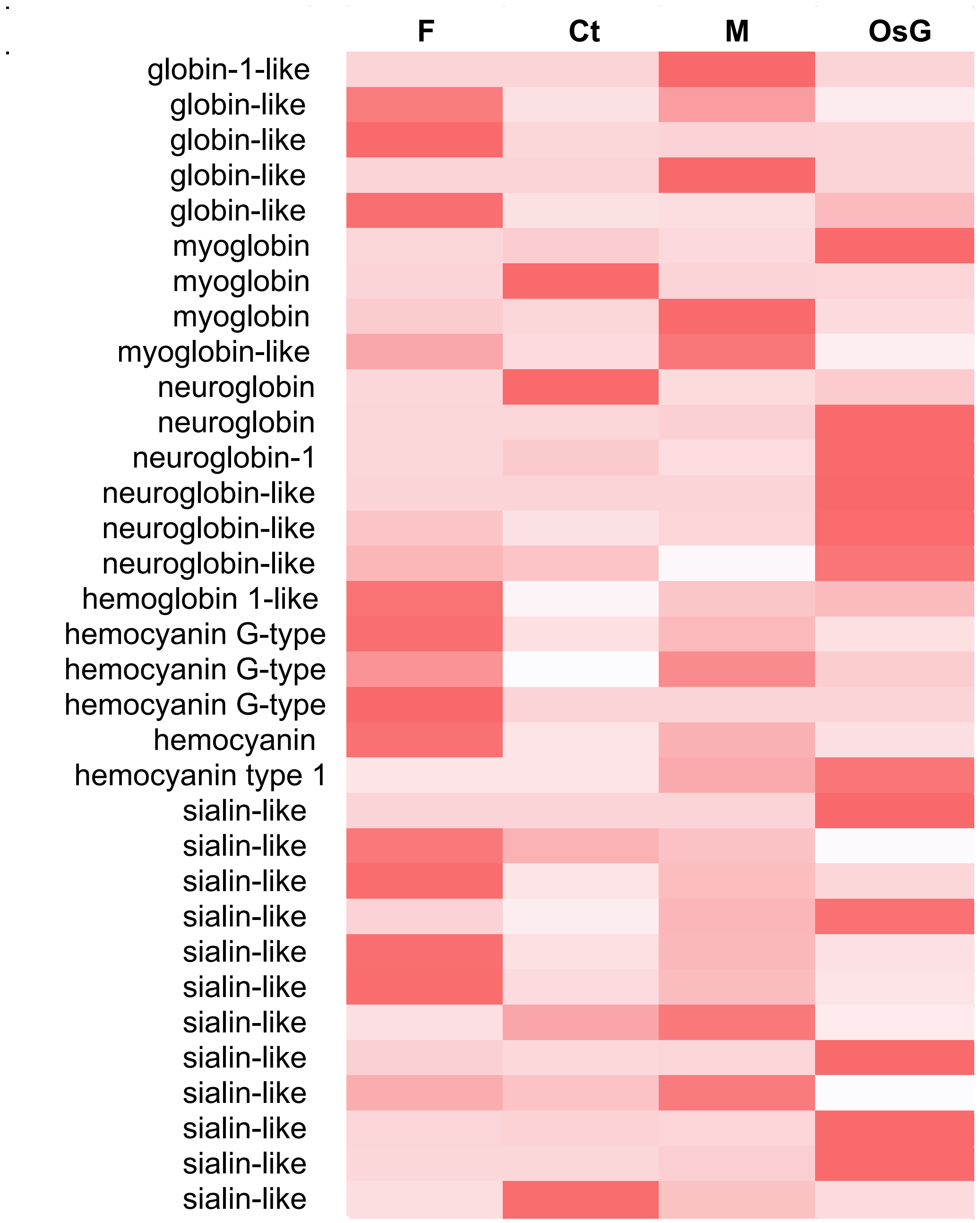


**Figure S11** A heat map showed the gene expression level of genes for transporting oxygen and nitrate (F: foot; Ct: ctenidium; M: mantle; OsG: oesophageal gland).

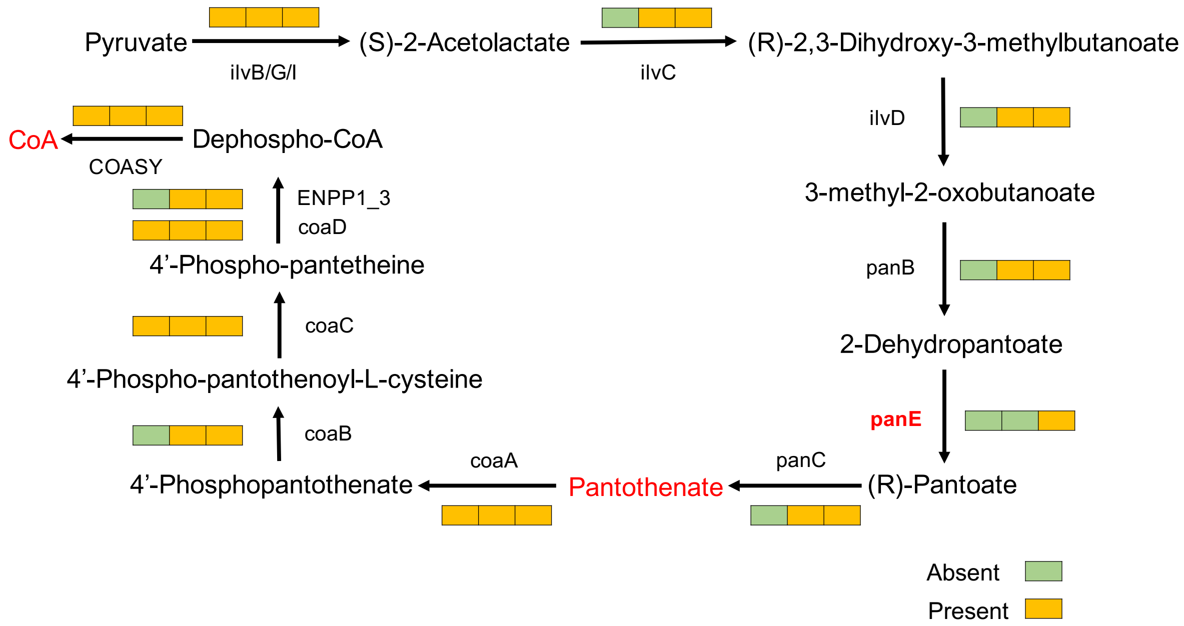


**Figure S12** Biosynthesis pathway of pantothenate (vitamin B5 in red) and coenzyme A (in red). The panE (in red) gene can only be found in the methane-oxidising endosymbiont. The block colour shows the absence (green) and presence (orange) of the gene in the genome. Left: host; middle: sulphur-oxidising endosymbiont, right: methane-oxidising endosymbiont.

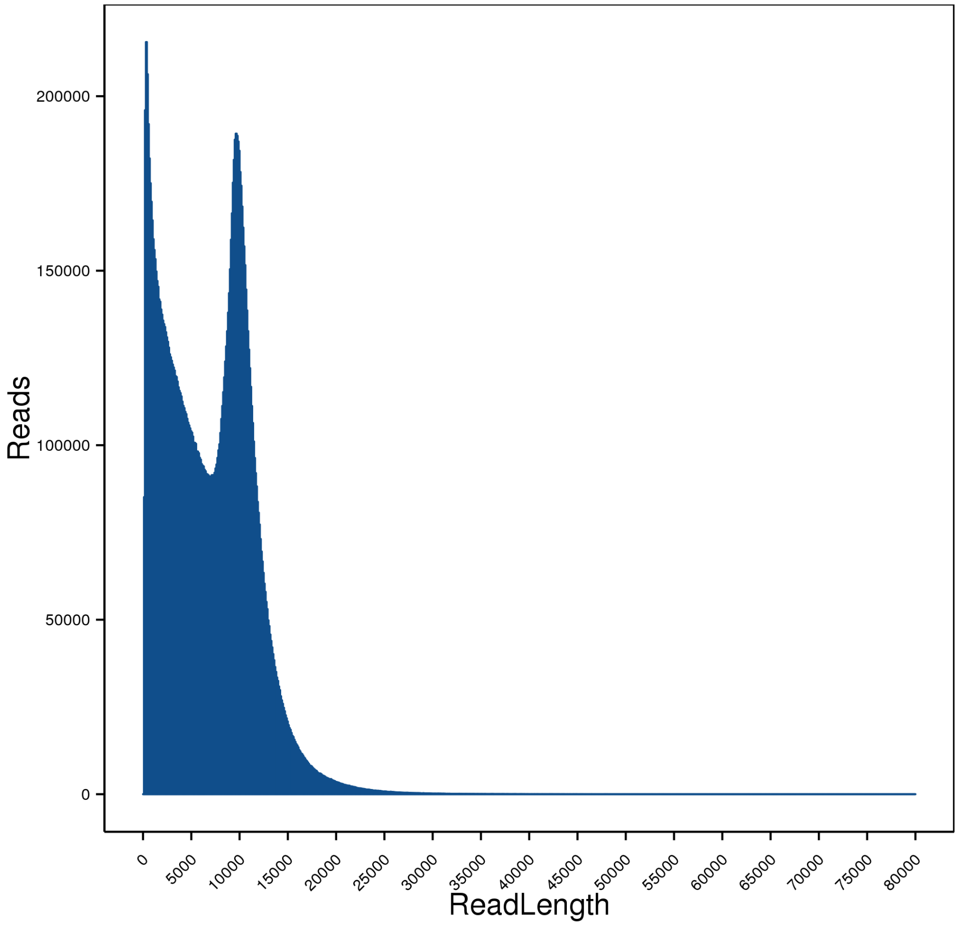


**Figure S13** The length distribution of PacBio raw sequencing subreads of *Gigantopelta aegis*.

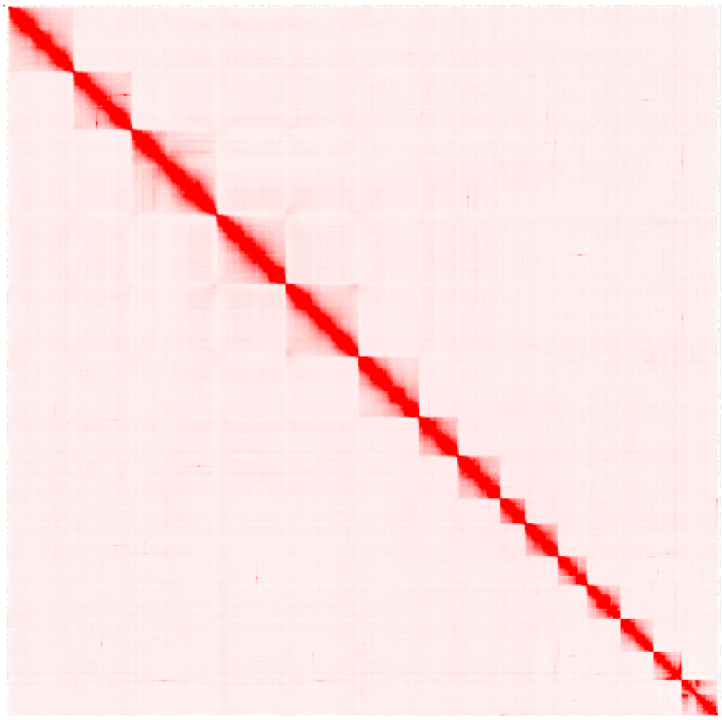

**Figure S14** The Hi-C contact map of 15 pseudo-chromosomes.

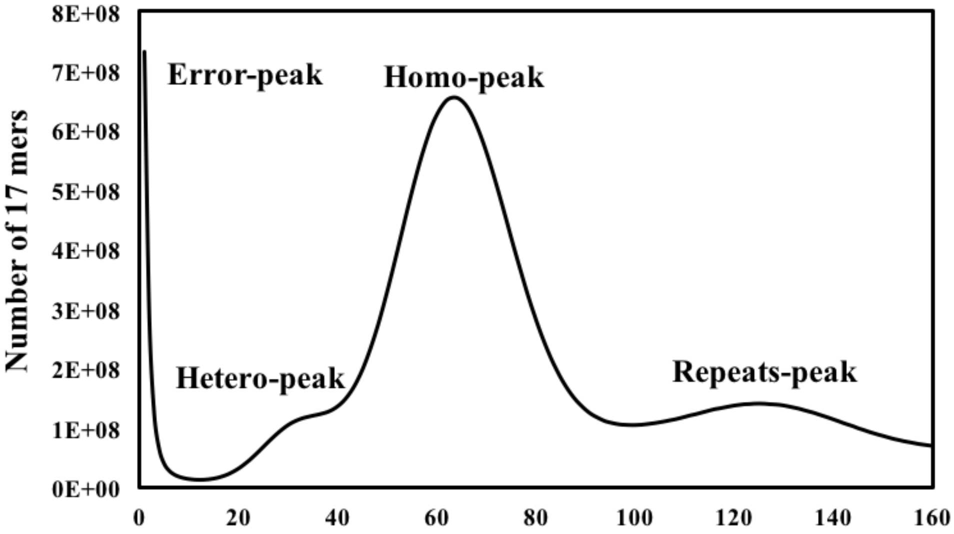


**Figure S15** The 17-mer distribution histogram of *Gigantopelta aegis* genome.

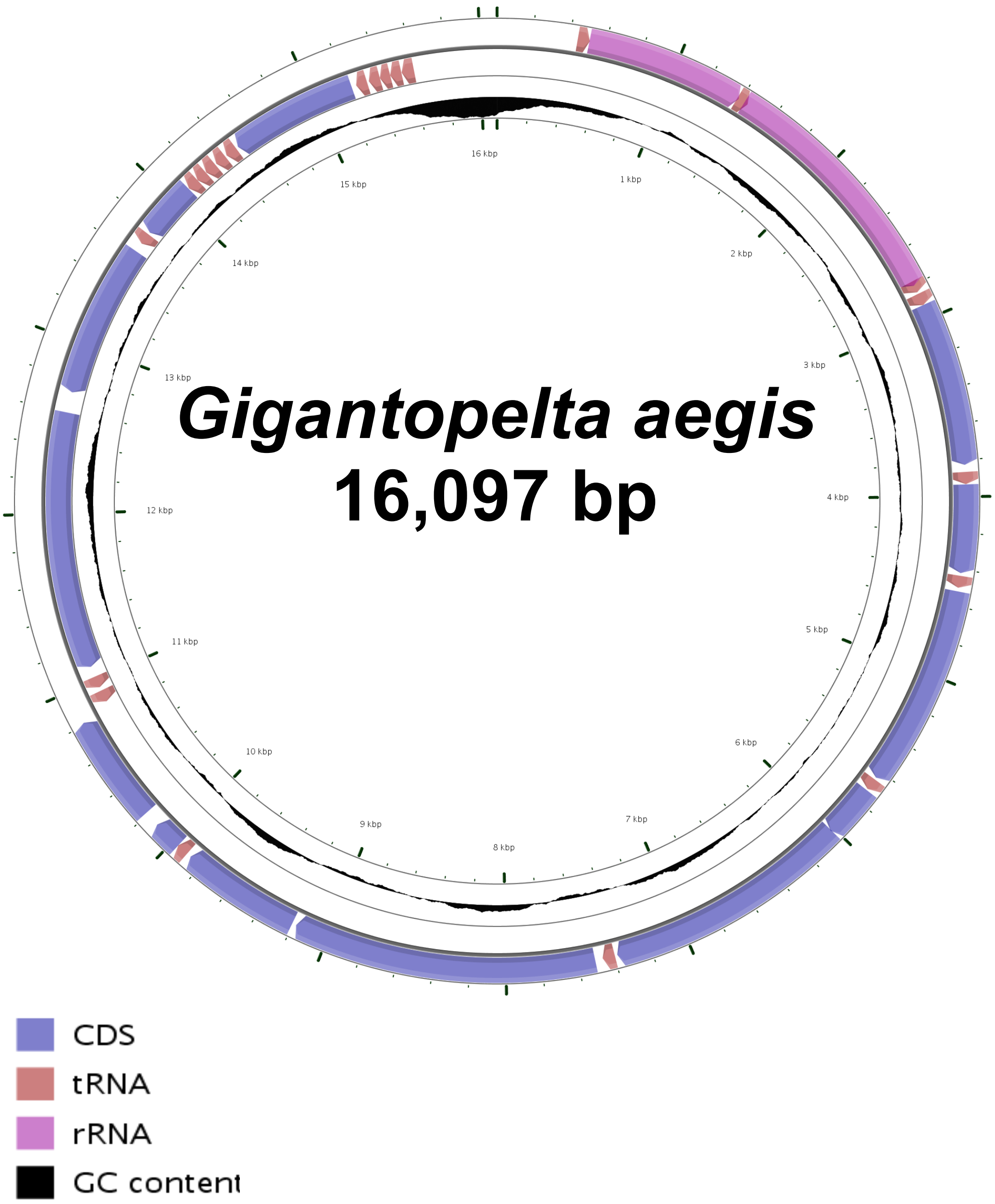


**Figure S16** The circos plot of the mitochondrial genome of *Gigantopelta aegis* (Inner ring: GC content; outer ring: genes order).

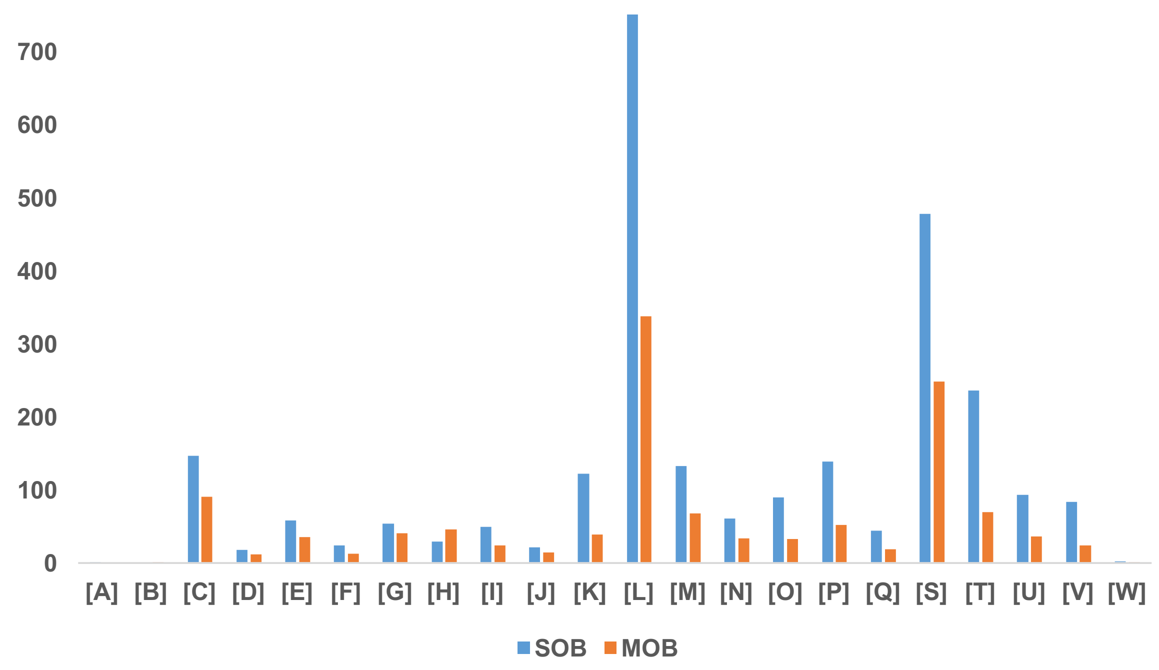


| | |
|---|---|
| [A] | RNA processing and modification |
| [B] | Chromatin structure and dynamics |
| [C] | Energy production and conversion |
| [D] | Cell cycle control, cell division, chromosome partitioning |
| [E] | Amino acid transport and metabolism |
| [F] | Nucleotide transport and metabolism |
| [G] | Carbohydrate transport and metabolism |
| [H] | Coenzyme transport and metabolism |
| [I] | Lipid transport and metabolism |
| [J] | Translation, ribosomal structure and biogenesis |
| [K] | Transcription |
| [L] | Replication, recombination and repair |
| [M] | Cell wall/membrane/envelope biogenesis |
| [N] | Cell motility |
| [O] | Post-translational modification, protein turnover, and chaperones |
| [P] | Inorganic ion transport and metabolism |
| [Q] | Secondary metabolites biosynthesis, transport, and catabolism |
| [S] | Function unknown |
| [T] | Signal transduction mechanisms |
| [U] | Intracellular trafficking, secretion, and vesicular transport |
| [V] | Defense mechanisms |
| [W] | Extracellular structures |

**Figure S17** The Clusters of Orthologous Groups (COG) annotation of the sulphur-oxidising bacteria (SOB: blue columns) and methane-oxidising bacteria (MOB: orange columns). One-letter abbreviation for the functional category was determined by the COG database.

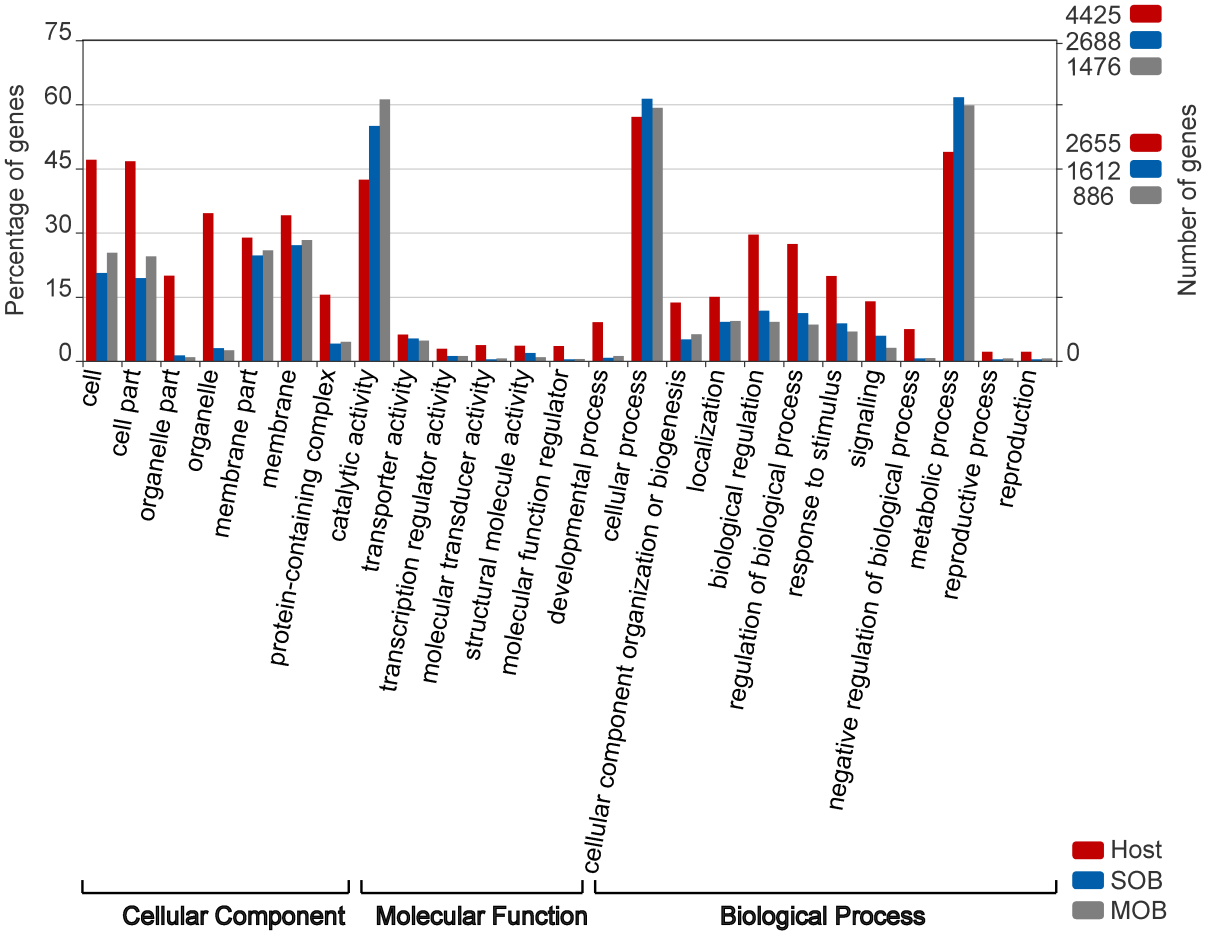


**Figure S18** A distribution plot shows the Gene Ontology (GO) items of the *Gigantopelta aegis* host, the sulphur-oxidising symbionts (SOB), and the methane-oxidising symbionts (MOB).

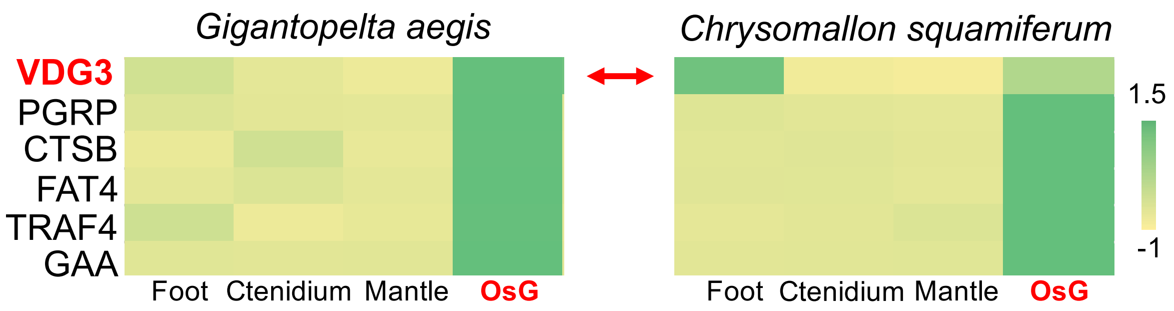


**Figure S19** The synteny blocks contain genes highly expressed in the oesophageal gland (OsG) of both *Gigantopelta aegis* and *Chrysomallon squamiferum*.
